# Episomal virus maintenance enables bacterial population recovery from infection and virus-bacterial coexistence

**DOI:** 10.1101/2024.07.30.605759

**Authors:** Rodrigo Sanchez-Martinez, Akash Arani, Mart Krupovic, Joshua S. Weitz, Fernando Santos, Josefa Anton

**Affiliations:** Department of Physiology, Genetics and Microbiology, University of Alicante, San Vicent del Raspeig, Alicante, Spain; Department of Biology, University of Maryland, College Park, MD, USA; Institut Pasteur, Université Paris Cité, Archaeal Virology Unit, Paris, France; Department of Physics, University of Maryland, College Park, MD, USA; Institut de Biologie, École Normale Supérieure, Paris, France; Multidisciplinary Institute of Environmental Studies Ramon Margalef, Alicante, Spain; Institute of Health and Biomedical Research of Alicante (ISABIAL), Alicante, Spain

**Author notes:** These authors contributed equally (Rodrigo Sanchez-Martinez, Akash Arani).

## Abstract

Hypersaline environments harbor the highest concentrations of virus-like particles (VLPs) reported for aquatic ecosystems. The substantial densities of both microbial populations and VLPs challenge traditional explanations of top-down control exerted by viruses. At close to saturation salinities, prokaryotic populations are dominated by *Archaea* and the bacterial clade *Salinibacter*. In this work we examine the episomal maintenance of a virus within a *Salinibacter ruber* host. We found that infected cultures of *Sal. ruber* M1 developed a population-level resistance and underwent systematic and reproducible recovery post infection that was counter-intuitively dependent on the multiplicity of infection (MOI), where higher MOI led to better host outcomes. Furthermore, we developed a nonlinear population dynamics model that successfully reproduced the qualitative features of the recovery. This suggests that the maintenance of the virus episomally, often referred to as pseudolysogeny, and lysis inhibition allow for host-virus co-existence under high MOI infections. Our results emphasize the ecological importance of exploring a spectrum of viral infection strategies beyond the conventional binary of lysis or lysogeny.

## INTRODUCTION

Diverse microorganisms adapted to life at high salt concentrations are found in globally distributed hypersaline environments that comprise ∼50% of continental waters (1). The abundance of prokaryotes increases with salinity, reaching values even higher than 10^8^ cells per milliliter at salinities above 25% (2, 3, 4). The microbial communities in these environments, especially at extreme salinities, are dominated by members of the domain *Archaea* along with *Bacteria* predominantly from the *Salinibacter* clade (5, 6). Amongst bacterial clades, *Salinibacter ruber* was the first bacterium confirmed to grow actively in extreme salinities (7). Global-scale biogeographic studies have revealed that *Sal. ruber* is one of the most prevalent and dispersed bacterial lineages in hypersaline environments. Moreover, it is also characterized by a high intraspecific diversity, which can be caused as a response to an intense viral predation (8, 9, 10). At saturated salt conditions, hypersaline environments are also characterized by the presence of viral-like particles (VLP) that can reach up to 10^10^ VLP/ml (11), more than three orders of magnitude higher than estimates from surface marine communities (12, 13). Measured ratios of viruses to microbes increase with salinity (from seawater to highly salt-saturated ponds) with values between 10 and 100 VLPs per cell, and can even reach 300 VLPs per cell (14). Hypersaline environments at close-to-saturation salinities are typically inhospitable to bacterivorous organisms found in other aquatic habitats, including protists and heterotrophic nanoflagellates (15, 16, 17). Unusually high viral abundances and virus-to-cell ratios along with reduced bacterivory in hypersaline environments suggest that viral infections and the subsequent modulation of microbial cell fate have a significant role in shaping prokaryotic population dynamics (18).

At the cellular scale, viral infections can lead to a continuum of outcomes spanning pathways that lead to rapid lysis as well as long-term persistence via lysogeny (19). In the lytic pathway, the virus redirects host metabolism to replicate the viral genome, produce virions, and lyse the host cell, thereby releasing viral progeny to the environment. In the lysogenic pathway, the infecting viral genome replicates along with the host chromosome. The provirus (i.e., the integrated viral genome) can reactivate stochastically or in response to changes in host state, re-initiating the lytic pathway leading to the release of virus particles (20, 21). Viral life cycles span the continuum between these archetypes, including life cycles such as chronic infections, in which there is a progressive release of virions into the environment without cell lysis (22). Another alternative viral life cycle is pseudolysogeny, in which genomes of non-temperate viruses persist inside host cells without integration as episomes, do not replicate synchronously with their host cell, and are transmitted asymmetrically from mother to daughter cells (23, 24).

The presence of high microbial densities, low bacterivory, and unusually high VLP levels in saturated brines presents a challenge to conventional explanations of top-down control. If extremely high viral densities are achieved through efficient infection and lysis, one might expect strong selection for viral-resistance via Kill-the-Winner like mechanisms (25, 26) or even the local collapse of archaeal and bacterial communities (and their viruses) until recolonized via dispersal. Alternatively, prokaryotes and viruses may persist at high levels in hypersaline environments because of sustained production from infected cells or a high stability of non-infectious viruses. Here, we utilize a *Sal. ruber*-virus model system to explore the mechanistic basis for the persistence and coexistence of high-density populations of viruses and bacterial hosts in hypersaline environments. Initial experiments revealed that virus infections led to a transient bacterial population crash followed by a recovery. However, contrary to our expectations and leveraging a combination of experiments, sequencing, and mathematical modeling, we conclude that bacterial population recovery was not enabled by the emergence of virus-resistant bacterial mutants or lysogeny. Instead, bacterial population recovery was enabled via the initiation of a persistent infection characterized by the episomal maintenance of viruses that protected infected cells from subsequent infection and lysis. As we show, this persistent infection enables lytic viruses to persist at high abundances without necessarily eliminating their hosts, providing new challenges to paradigms of virus-microbe coexistence in extreme environments.

## RESULTS

### Systematic and reproducible recovery of *Sal. ruber* populations after viral-induced lysis

We infected *Sal. ruber* strain M1 with the virulent M1EM-1 virus (27) (named henceforth EM1 for convenience) at a MOI=0.01 in triplicate at time=0 h (Fig. 1a). In all three replicates, we observed a rapid stabilization in the host optical density (OD) following infection due to lysis, followed by a population-level recovery at 80 h such that infected cultures eventually recovered to the levels comparable to those of the virus-free control, reaching the stationary state ODs (see Methods for further details). This systematic recovery of the population post infection was also observed in other *Sal. ruber*-virus pairs (see Extended Data Fig. 1). This finding was unexpected as the EM1 virus does not encode lysogenic markers (i.e., excisionases and integrases) which would allow its integration into the host chromosome, facilitating superinfection exclusion-mediated population level recovery as seen for some temperate viruses (28). We further examined this recovery phenomenon by infecting *Sal. ruber* M1 with the EM1 virus at ten-fold higher viral levels (MOI=0.1) and using more replicates (10 replicates for both control and infection treatment; Fig. 1b). In this case, we observed a sharp decrease in the OD levels of the infected cultures, consistent with the 10x increase in viral pressure (MOI=0.1 vs. MOI=0.01). However, we once again observed recovery dynamics comparable in all ten replicates.

**Fig. 1:**
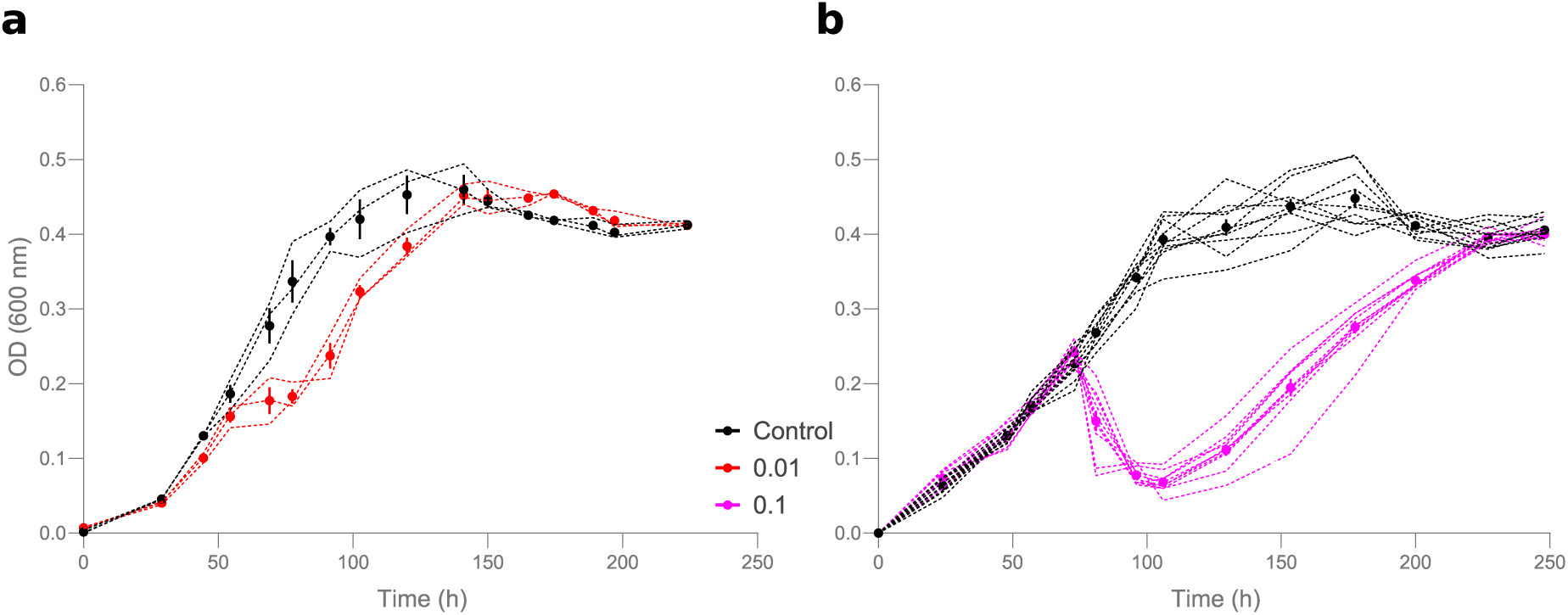
Infection experiments show a reproducible recovery after viral lysis. Infection experiments of *Sal. ruber* strain M1 with the EM1 virus were performed using two different MOIs. Black lines represent uninfected control and red (**a**, MOI=0.01) and purple (**b**, MOI=0.1) lines represent virus-infected cultures. Each of the individual replicates used are shown as dotted lines. The dots show the mean and the vertical bars the standard error. The experiments were performed with 3 replicates for MOI 0.01 and 10 replicates for MOI 0.1 for both the control and infected cultures. Viruses were mixed with the host at time=0 h.

### *Sal. ruber* recovery is not consistent with the proliferation of virus-resistant mutants or the integration of viral genomes into bacterial genomes

We hypothesized that reproducible recovery of infected cell cultures is not caused by spontaneous mutations and the emergence and proliferation of virus-resistant bacterial populations which typically involves jackpot like variability that is incompatible with the observed reproducibility in recovery times (29). Nonetheless, to assess the link between recovery and virus-resistance, all replicates (both the infected cultures and the controls) from the recovered cultures shown in Fig. 1a (named C_F_) were sequenced. The initial *Sal. ruber* M1 used as a host (named C_0_) was also sequenced as a control. We then analyzed and compared the sequenced genomes to evaluate the evidence for selective sweeps of virus-resistance genes in infected replicates.

Only 6 mutations showed a frequency close to 100% (Supplementary Table 1) in all the infected replicates. Of these 6 mutations, 3 involved non-synonymous changes, affecting an ATP-dependent helicase/deoxyribonuclease, an orotate phosphoribosyltransferase and a sodium/glucose cotransporter. The first two are related to helicase/exonuclease activities and to the synthesis of pyrimidines, respectively, so they did not seem to have a direct relationship with virus resistance. The third mutation, affecting the stop codon in the synthesis of the sodium/glucose cotransporter, could be involved in the resistance, as it affects a transmembrane protein. However, upon further examination, we found that both resistant and sensitive cells had the same mutation, confirming that it was unrelated to resistance. In addition, it was subsequently found that cells that regained susceptibility to the virus retained all the mutations mentioned above (see below and *Mutations Supplementary Information* for further details). Therefore, no mutations could be unequivocally tied to the observed population recovery phenomenon in the infected cultures.

According to previous studies (30), these dynamics after viral infection could be explained by the integration of the viral genome into the chromosome of its host, yielding lysogens resistant to superinfection with the same virus. Although EM1 lacked any lysogeny-associated traits in its genome, including excisionase and integrases (Supplementary Table 2), the reads of the infected C_F_ cultures were reexamined to search for hybrid host-virus reads, which would signify integration into the chromosome (see Methods). No hybrid reads were found, and therefore recovery cannot be attributed to virus integration. However, we observed a remarkably high sequencing depth of the virus genome within infected C_F_ cultures, averaging 146 nt/position, while the cellular chromosome exhibited 116 nt/position (Supplementary Table 3). This similar level of sequencing depths suggests that viral genomes are maintained intracellularly in the recovered cultures, albeit not via integration as a prophage.

### Infected cells with episomally maintained viruses comprise part of the recovered *Sal. ruber* population

The lack of evidence for mutations related to virus resistance or integrated prophage within recovered *Sal. ruber* populations led us to hypothesize that recovered bacterial populations infected by viruses might maintain viral genomes intracellularly in an episomal form. To explore this alternative, we grew colonies from time point C_0_ (initial *Sal. ruber* M1) and C_F_ (final recovered cells). Twenty individual colonies from both C_0_ and C_F_ were grown in liquid medium and PCR-tested for the presence of the EM1 virus. As expected, all C_0_ colonies tested positive with the specific primers for *Sal. ruber* M1 but tested negative for the virus. However, all the final colonies from the infected cultures tested positive for the EM1 virus. Three randomly selected cultures from the C_F_, colonies that tested PCR positive for the virus (which we refer to as 1R, 2R and 3R respectively), were stained with SYBR^TM^Gold and observed under the epifluorescence microscope (Fig. 2a). After observation of more than thirty fields per culture, extracellular virions were not observed in any case. Hence, although EM1 virus was present in the culture, its virions were not detected extracellularly. The intracellular presence of viral genomes was confirmed by pulsed field gel electrophoresis (PFGE) with DNA from the final colony 1R, but not from the control C_0_ colony (wild type *Sal. ruber* M1). The PFGE showed a band with the size of the EM1 virus genome (35 kb) in 1R, which was confirmed by Southern blot hybridization with an EM1-specific probe (Fig. 2b). These results, in conjunction with the sequencing data that revealed no signs of integration (no hybrid host-virus reads were found), the lack of integrases and/or recombinases in the genome of EM1 (Supplementary Table 2), supports the hypothesis that the EM1 virus was maintained in an episomal form within *Sal. ruber* M1 recovered populations. In addition, the terminal redundancy found in sequencing may indicate that it is maintained within cells as a circular episome.

**Fig. 2:**
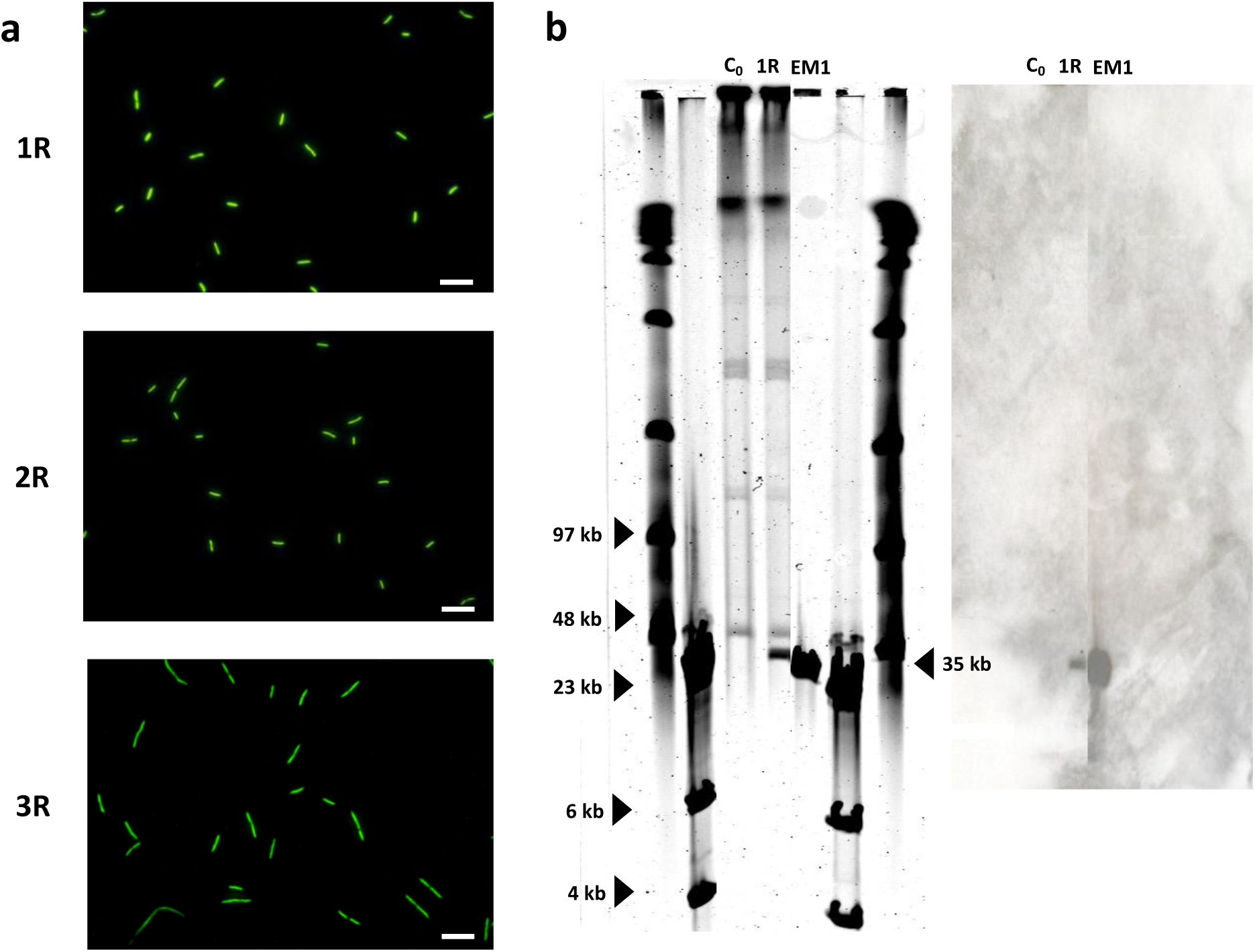
The EM1 virus establishes a pseudolysogenic state in *Sal. ruber* M1. **a**, Three randomly selected cultures of the 20 colonies of C_F_, PCR positive for the EM1 virus, that we named 1R, 2R and 3R, were stained with SYBR^TM^Gold and observed under epifluorescence microscope to test the presence of extracellular viruses. No extracellular viruses were seen in any of them. 30 fields were examined per sample. The photos shown here are a representation of each sample. Scale bar: 10 μm. **b**, A PFGE was performed (left) comparing initial cells (C_0_) with final colony 1R (lines 3 and 4 of the gel, respectively). The DNA of the virus was also loaded as a control (line 5). The PFGE showed a band in 1R with the size of EM1 (35kb) which was not found in C_0_. A southern blot (right) with a labeled probe targeting the genome of EM1 confirmed that this band belonged to the virus.

### Episomal maintenance of viruses protects recovered cells from subsequent infection and viral-induced lysis

To test whether the acquisition and episomal maintenance of the virus had an influence on the recovered cell growth, we conducted growth experiments on 1R and compared the population dynamics to that of the wild type *Sal. ruber* M1 (Fig. 3a). We found no statistical differences in the reached OD or growth dynamics between 1R and *Sal. ruber* M1 (*Modeling Supplementary Information: Sal. ruber M1 wild-type and 1R (pseudolysogen) growth comparison*). Hence, there is no evidence that viral acquisition provides a direct cost or benefit to host growth. However, the benefits of intracellular maintenance of viral genomes may be contextual.

**Fig. 3:**
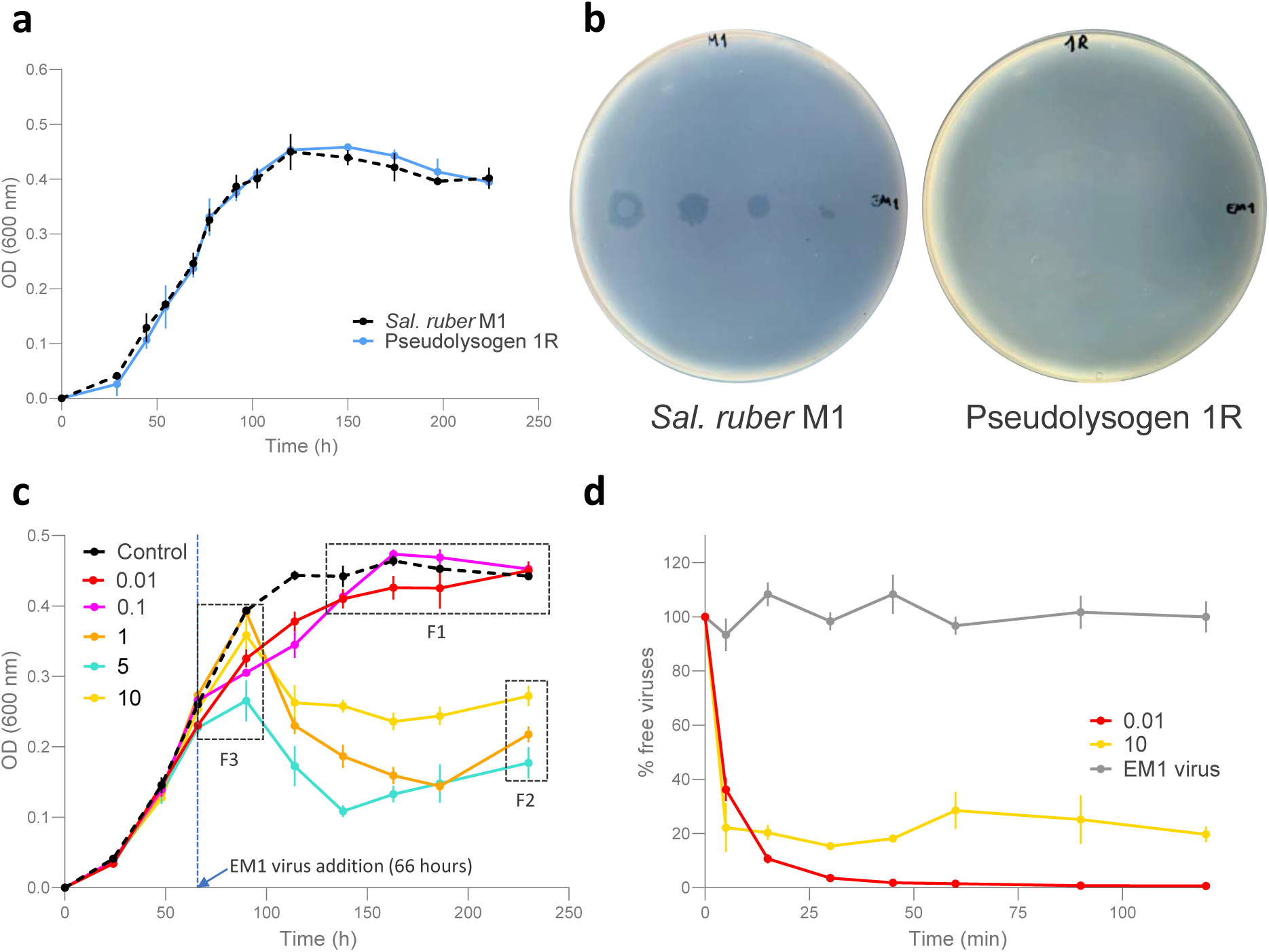
Viral acquisition protects Sal. ruber M1 from further infection and has MOI dependent properties. **a**, Growth curves (performed in triplicate) of *Sal. ruber* M1 wild type (black dotted line) and pseudolysogen 1R (blue line) were performed and their growth dynamics were found to have no statistical differences (*Modeling Supplementary Information: Sal. ruber M1 wild-type and 1R (pseudolysogen) growth comparison*). **b**, Susceptibility to the virus was assessed using a spot test. *Sal. ruber* M1 wild type (left) and pseudolysogen 1R (right) were exposed to 3 μl of the EM1 virus tittered at 10^10^ PFUs/ml (left part of each plate) and diluted 10^2^, 10^4^ and 10^6^ (left to right) times. **c**, Infection curves of *Sal. ruber* M1 and the EM1 virus were performed at different MOI. The M1 cultures were infected at 66 hours, during the exponential growth phase. The black dotted line represents the non-infected control and the different colors indicate the MOI, as is indicated in the legend. The experiment results exhibit three key features (*see text).* **d**, An adsorption assay of the EM1 virus to *Sal. ruber* M1 wild type at MOI 0.01 (red line) and 10 (yellow line), where the number of extracellular viruses (PFU/ml) was measured over 120 minutes. The virus stock used was also set as a decay control (grey line). All the experiments were performed by triplicate. Error bars represent the standard error.

During typical lysogeny, the prophages can protect hosts from viral infection and lysis via superinfection exclusion (31, 32, 33). We hypothesize that similar principles may apply to recovered bacterial populations. To test for superinfection exclusion in recovered populations, we conducted spot test assays with 1R and the wild type *Sal. ruber* M1. In both cases, we plated cells and then overlaid agar plates with spots of EM1 virus serially diluted from 10^10^ PFU/ml to 10^4^ PFU/ml (see Methods). The wild type strain was susceptible to EM1 infection, whereas 1R exhibited resistance (Fig. 3b). We interpret the results of the spot assay experiment to imply that episomal maintenance of the EM1 virus confers protection against subsequent EM1 infection and lysis. Furthermore, by conducting an adsorption assay with 1R and the wild type *Sal. ruber* M1, we found that protection against subsequent EM1 infection occurs at the surface level, i.e., preventing effective adsorption (Extended Data Fig. 2).

Finally, to test how stable is the intracellular maintenance of EM1, we plated the 1R strain and picked colonies at random to test for the presence of the virus by PCR. The results showed that 13 of 17 colonies retained the virus (*Mutations Supplementary Information*). A spot test with EM1 on the cultures derived from the 17 colonies revealed that only those strains that had lost the virus genome were sensitive to EM1 infection, confirming that the resistance was due to the virus acquisition and episomal maintenance. Subsequent genomic sequencing of 3 of these sensitive and 3 resistant colonies (all derived from 1R) showed that the mutations observed in recovered cultures following the infection (see above) had not been lost and therefore were not involved in the development of resistance to EM1 infection (*Mutations Supplementary Information*). The reappearance of sensitive strains within the 1R recovered population suggests that the viral genome is not replicating synchronously or faithfully segregated with the host chromosome. Consistently, no marker was found in the virus genome that would indicate cell-synchronous replication and partition, such as the Par system of bacteriophage P1 (34) (Supplementary Table 2). Therefore, we hypothesize the viral persistence inside recovered cells is enabled through asymmetric episomal virus transmission during cell division consistent with a pseudolysogenic state (see (24)). For the remainder of the manuscript, we refer to bacterial populations post-infection as ‘recovered’ and use the term resistant pseudolysogens to refer to the subset of bacterial populations that have episomally maintained the virus genome and are resistant to subsequent infection and lysis by the same virus.

### Bacterial population recovery depends non-monotonically on MOI

The spot assay experiment yielded a counter-intuitive result. When high virus concentrations were added on the plates of wild type *Sal. ruber* M1 at levels between MOI of 10-100, we observed growth of cells in the center of spots. This suggests that there may be a MOI dependence to the initiation of pseudolysogeny, akin to that found in classic studies of lysogeny in which the probability of lysogeny increases with increasing MOI (21, 35, 36). Hence, cultures with cells infected by multiple viruses may be more likely to yield pseudolysogens which can grow even in the presence of viruses. To explore the MOI-dependence of recovery further, *Sal. ruber* M1 was infected with EM1 at MOIs ranging from 0.01 to 10 viruses per host cell. The M1 population was infected during its exponential growth phase (66 h). The population dynamics (Fig. 3c) exhibit dip and recovery dynamics for MOI 1, 5 and 10 in a similar fashion to the initial infection curve experiment. In contrast, population dynamics for MOI 0.01 and 0.1 solely exhibit growth, albeit slightly slower than in the control population. Notably, the final cell densities in the experiments with MOI 1, 5 and 10 were non-monotonically related to MOI, i.e., the OD values were 0.22 +/- 0.01, 0.18 +/- 0.02 and 0.27 +/- 0.01 for MOI 1, 5, and 10, respectively. The OD value at time 230 hours for MOI 10 was statistically significantly different from the OD value for MOI 1 (t-test, p=0.041) and MOI 5 (t-test, p=0.038). Note that we could not reject the null hypothesis that the OD values at time 230 hours for MOI 1 vs 5 were statistically different (t-test, p=0.206). These results raise the question: if EM1 generates resistant pseudolysogens (as was confirmed in the spot assay experiment for 1R (see Fig. 3b)), then why would recovery dynamics depend non-monotonically on MOI?

Typically, superinfection exclusion requires certain cellular rearrangement post-infection (31, 32, 37). As such, we hypothesize that multiple EM1 viruses can jointly infect the same *Sal. ruber* M1 cell, insofar as multiple infections precede the development of resistance (38). For example, given a MOI of 10 and assuming that all virions have adsorbed to the cell and injected their genomes, the infection would yield an average of 10 viruses per cell with a standard deviation of ∼3. Therefore, to test whether superinfection was possible in this system, we performed adsorption assays with the wild type *Sal. ruber* M1 and EM1 at MOI of 0.01 and 10 (Fig. 3d). In the case of MOI 0.01, the decrease of free viruses was equivalent to that expected in theory for complete adsorption of virions onto cells (i.e., approximately 1% of cells should be infected). In the case of MOI 10, the exponential decrease of free viruses plateaus at ∼20%, implying that 80% of virions adsorbed to cells, consistent both with superinfection (i.e., the average cellular-level MOI would be 8) and interference that limits viral superinfection at high MOI. Hence, we conclude that superinfection of *Sal. ruber* M1 is possible by the EM1 virus given sufficiently short intervals between infection events even if the development of a superinfection exclusion state prevents subsequent infection over the long term in resistant pseudolysogens.

### Scaling-up the dynamical effects of pseudolysogeny from cells to populations

The contextual impact of superinfection raises questions on how pseudolysogeny modulates recovery dynamics, in light of three key empirical observations (Fig. 3c). First, the *Sal. ruber* M1 populations infected at MOI 0.1 and 0.01 do not exhibit an observable population decline. Second, the M1 host population recovers to a higher population densities when infected at MOI 10 compared to MOI 5 and MOI 1, highlighting a non-monotonic relationship between viral levels and population-level lysis. Third, when infected at MOI 10, the *Sal. ruber* M1 population does not crash immediately upon infection and maintains persistent transient growth post infection. To connect mechanisms occurring at cellular-scales with these population-level dynamics, we developed a nonlinear ordinary differential equation (ODE) model of the EM1 virus interacting with *Sal. ruber* M1 hosts (Fig. 4). The model includes dynamics of resources (*R*), sensitive cells (*S*), pseudolysogens (*P*) and virus particles (*V*) (39). We separate pseudolysogens into “early” (*P_e_*) and “fully developed” pseudolysogens (*P_f_*), where viruses can infect sensitive cells and early pseudolysogens (consistent with findings of Fig. 3d) which then transition to become resistant, *P_f_* types (consistent with findings of Fig. 3b). We account for cellular-level multiplicity of infection of the pseudolysogens through a superscript ^[*k*]^ where *k* denotes the number of viral genomes per cell e.g., *P_e_^[2]^* denotes the number of early pseudolysogens with 2 viral genome copies. The number of viruses able to infect a single cell is limited, consistent with adsorption results from Fig. 3d (details in Extended Data Fig. 3). Cell division for both sensitive cells and pseudolysogens is modeled as a function of resource consumption. Full details of the model specification and parameters are provided in the Supplement (*Modeling Supplementary Information: Main Model)*.

**Fig. 4:**
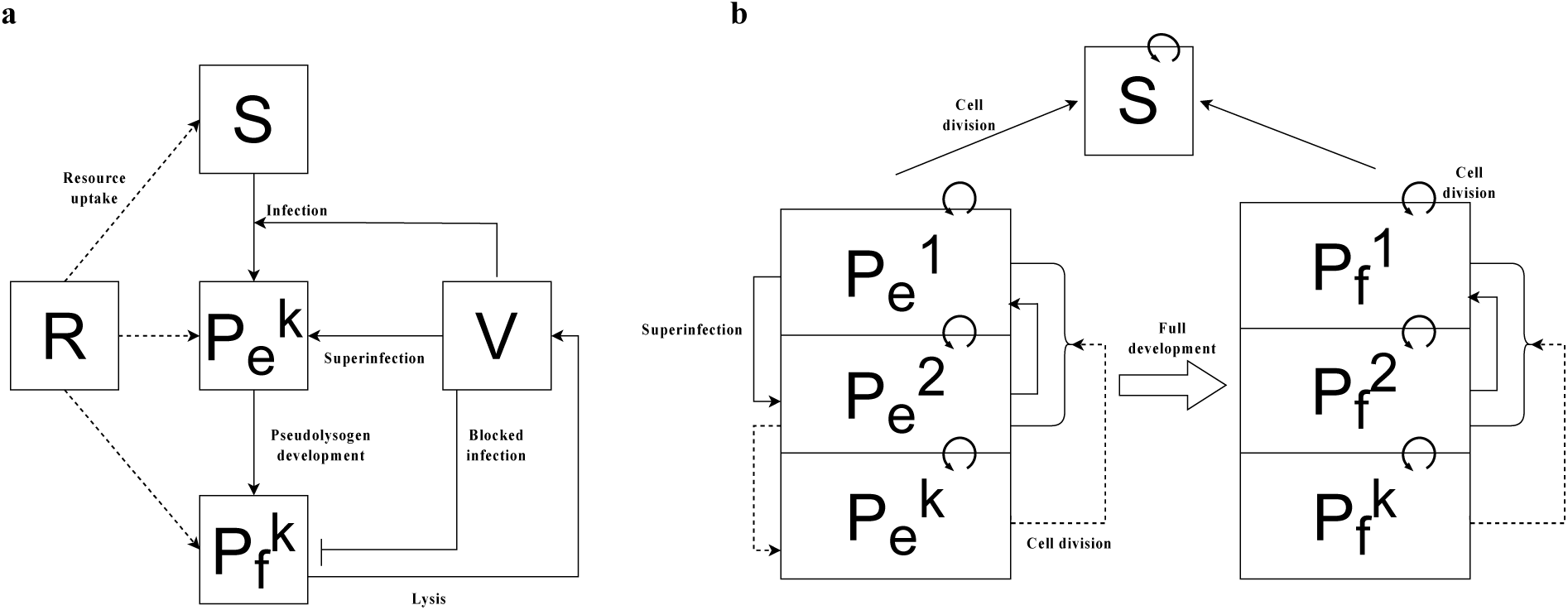
Schematic of nonlinear dynamics population model. **a,** Schematic representation of the ODE model. *R* denotes the resource level and *V* denotes the free virus concentration. The host population is split into three classes: sensitive (*S*), early pseudolysogen (*P_e_^k^*), and fully developed pseudolysogen (*P_f_^k^*), where *k* is the number of viral genome copies within the host cell. **b,** Dynamics of pseudolysogen growth and superinfection assuming primary infection has already occurred (primary infection: *S → P_e_^1^*). Early pseudolysogens get superinfected causing *P_e_^k^* → *P*_*e*_^*k*+1^transitions, while fully developed pseudolysogens are resistant to superinfection at the surface level (denoted by *blocked infection* arrow). Pseudolysogen cell division for both early and fully developed pseudolysogens randomly splits the viral genome of the parent, causing *P*^*k*^ → *P*^*k*′^transitions where *k’* is a randomly chosen positive integer that is *≤ k* (*P^0^* is virus-free and is therefore equivalent to *S*). Early stage pseudolysogens eventually transition into fully developed pseudolysogens that still cell divide but are resistant to superinfection.

Simulated population dynamics (Fig. 5a) recapitulate the key empirical observations (as shown in Fig. 3c). The match between theory and experiment is driven by the following mechanisms. First, the model assumes that host lysis decreases with decreasing resources (40, 41). In Fig. 5b, we show that decoupling host lysis from resource concentrations leads to a rapid crash in *Sal. ruber* dynamics amongst the low MOI populations that is inconsistent with experimental observations. Second, the model assumes that cellular lysis rates decrease as a function of increasing cellular MOI (42, 43). In Fig. 5c, we show that if lysis rates are independent of cellular MOI, then population crashes are most apparent at the highest MOI and decrease with decreasing MOI, inconsistent with experimental observations. Third, we assume that pseudolysogens divide and that their daughter cells can inherit viral genomes, albeit randomly, from the parent cell (akin to random segregation of episomal elements (44)). In Fig. 5d, we show that if pseudolysogens do not divide then the *Sal. ruber* population at MOI 10 (yellow curve) immediately dips upon infection at 66 hours (blue dotted line) and there is no persistent transient growth, thereby inconsistent with the experimental dynamics. In *Modeling Supplementary Information*: *Subsets of main model with corresponding dynamics,* we explore all potential combinations of the presence/absence of each mechanism and find that the model only recapitulates observed dynamics when all three mechanisms operate jointly. Hence, we contend that each of these elements (resource-dependent lysis, latent period delays in multiply infected cells, and pseudolysogen cellular division) are necessary to explain experimentally observed dynamics of the EM1 virus and its host across 3 orders of magnitude difference in initial ratios of virus to host. The details of how all the model features are implemented is highlighted under *Modeling Supplementary Information: Key Model Features* and *Modeling Supplementary Information: Viral passage in superinfected pseudolysogens*. Finally, we note that the results are robust to variation in the variability associated with the passage of episomally maintained viruses to daughter cells during replication (*Modeling Supplementary Information: Episomal viral replication*).

**Fig. 5:**
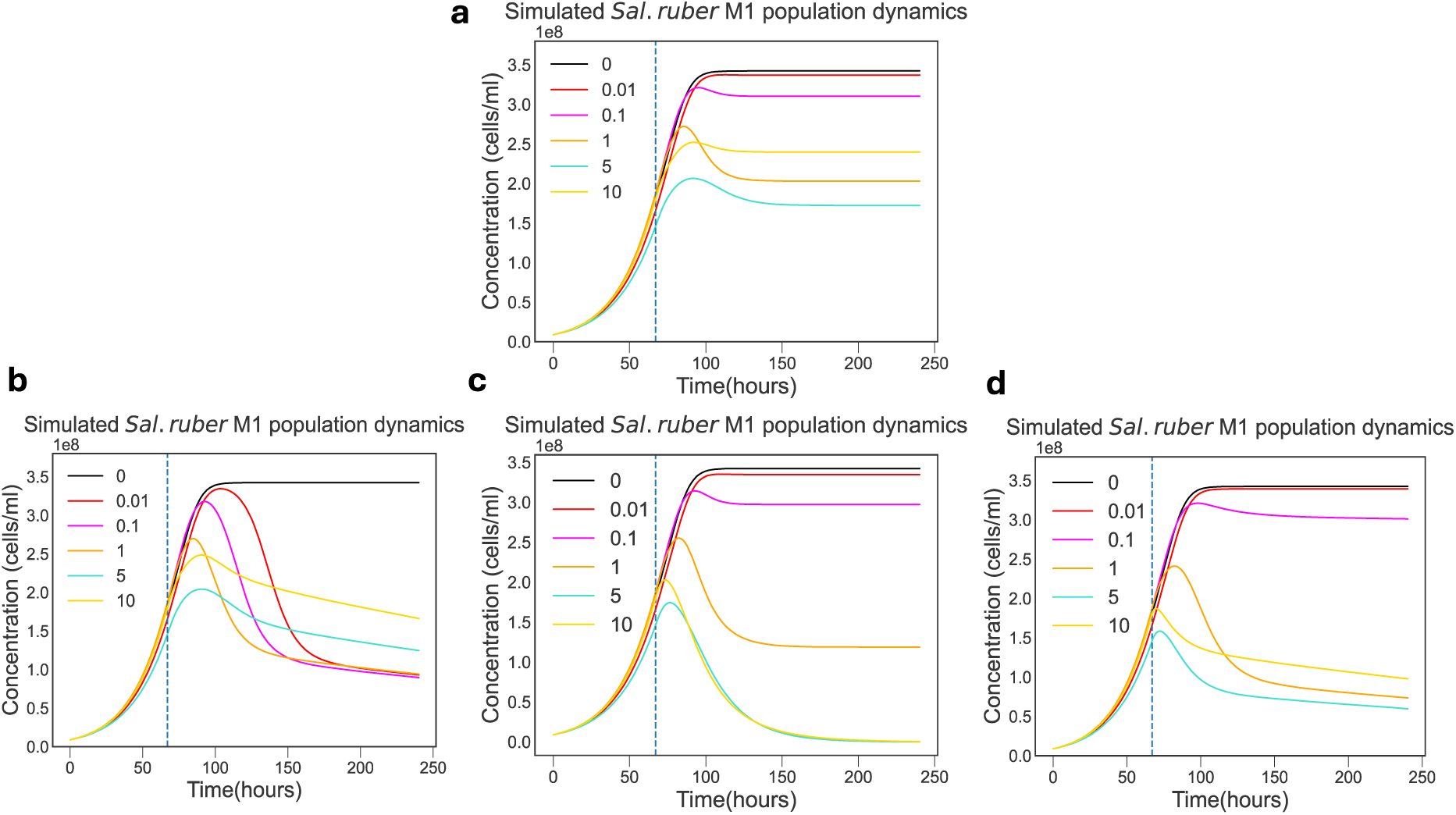
Simulated population dynamics recapitulate experimental features through implementation of lysis inhibition and pseudolysogenic cell division. **a,** Simulated host population dynamics 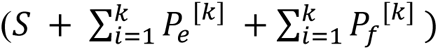 using the ODE model proposed *(Modeling Supplementary Information: Main Model)* to successfully recapitulate the experimental features highlighted in Fig 3c. Subfigures b,c,d graphically show how decoupling features from the model prevents simulations from capturing experimental features. **b,** Simulated host population dynamics with lysis rates independent of resource levels. **c**, Simulated host population dynamics with lysis rates independent of cellular MOI. **d**, Simulated host population dynamics without pseudolysogen cell division.

### Episomal maintenance of viruses and the population-scale maintenance of virus-microbe coexistence

Finally, we set out to explore the extent to which episomal maintenance of viruses inside recovered bacterial populations could enable virus-microbe coexistence over longer time scales. The persistence of the virus within the pseudolysogen population was tested by transferring pseudolysogen 1R every 9-10 generations to fresh medium. We made four such transfers, resulting in ∼ 40 generations of pseudolysogen growth and cell division (Extended Data Fig. 4a). At the end of the exponential phase of each transfer, we conducted a PCR test and found that the virus was not lost over time (Extended Data Fig. 4b). Upon qPCR analysis, we found that the ratio of virus genome/chromosome was close to 1 in the first transfer and upon further transfers, the ratio increased to 2-3 virus genomes per chromosome of M1 and was maintained at this level (Extended Data Fig. 4c). To determine whether viral resistance was maintained within the population, cells from each of the aforementioned transfers were exposed to EM1 through a spot test. We found that the cells from every transfer exhibited resistance to additional infection (Extended Data Fig. 5). Next, to observe how the pseudolysogens might ecologically interact with other *Sal. ruber* strains, we conducted spot tests in which eight strains of *Sal. ruber* used routinely in our laboratory were individually exposed to pseudolysogen 1R. We observed that *Sal. ruber* M1, the only strain sensitive to the EM1 virus, was killed while the rest of the seven strains were maintained (Extended Data Fig. 6). Similar dynamics have been observed with temperate viruses, where lysogens induce and release free virions as a weapon to kill sensitive cells in shared environments (45). We hypothesize that a subset of pseudolysogens of *Sal. ruber* reactivate the lytic pathway, releasing virions that infect sensitive competitors, generating a mix of pseudolysogens and increasing the level of extracellular virions.

## DISCUSSION

Here we explored the dynamics arising from interactions between a virulent virus (EM1) and one strain of a prevalent and globally dispersed bacterial host found in hypersaline environments (*Sal. ruber* M1). We found that *Sal. ruber* M1 populations reproducibly recover from infections by EM1 viruses across three orders of magnitude variation in MOI. Notably, M1 bacterial populations recover to higher densities when infected with more viruses (MOI=10) than with less viruses (MOI 1 or 5) while recovering to densities comparable to those of non-infected controls for MOI levels of 0.1 or 0.01. Both sequence and experimental evidence suggest that recovered populations have a significant fraction of cells with episomally maintained virus genomes that pass asymmetrically from mother to daughter cells in the absence of synchronized replication with their cellular hosts. Subsequent infection experiments revealed that these cells with episomally maintained virus genomes (i.e., pseudolysogens) are resistant to infection and lysis by the original virus, providing an opportunity to explore mechanisms by which infection recovery could stabilize bacteria-virus feedback. We developed and analyzed a nonlinear population dynamics model that recapitulated key features of the MOI dependent recovery dynamics that depend on the potential joint infections of cells, lysis inhibition via superinfection immunity, and lysis inhibition due to resource limitation. Altogether, the initiation and maintenance of pseudolysogeny stabilizes short-term population dynamics while providing a mechanism for long-term coexistence of viruses and their bacterial hosts in extreme environments.

The current study connects cellular-level interactions to feedback mechanisms that stabilize virus-host population dynamics that could otherwise lead to population collapse, especially at high MOI values. While previous studies have shown similar recovery dynamics in different virus-host systems (46, 47, 48, 49), there has been limited effort to explain the mechanisms responsible. Studies observing recovery dynamics phenomenon with temperate viruses have pointed to emergence of resistant lysogens which are protected against further infections (28, 30). However, bacterial population recovery has also been observed in viruses without any lysogeny markers as well (50, 51), in which the emergence of resistance has been overlooked or simply attributed to spontaneous mutations of the host (52, 53). Here, recovery dynamics could not be attributed to lysogeny given the use of the *Sal. ruber* M1-EM1 system where EM1 is strictly virulent. The recovery dynamics observed here was also highly reproducible, inconsistent with recovery due to the emergence and expansion of spontaneous resistant mutants that grow from low to high frequency where substantial variability is a hallmark feature of dynamics (29). Furthermore, our analysis revealed no clear resistance-associated mutations in the recovered M1 populations. Similarly, we also identified lysis inhibition at high MOI and/or when resources are limited as an essential part of explaining the MOI-dependence of recovery dynamics. Lysis inhibition has commonly been noted in infection of *Escherichia coli* by T4 (40, 42, 53, 54), with some studies suggesting that it plays a key role in the maintenance and development of *E. coli*-T4 pseudolysogens (24, 54). We hypothesize that lysis inhibition could similarly be a key feature of pseudolysogeny in the *Sal. ruber* M1-EM1 system, allowing for persistent infection in starved conditions and/or under high MOI.

Our combined study comes with caveats. First, we focused primarily on contextual benefits of pseudolysogeny, noting that identifying direct impacts of episomal maintenance of viruses on cellular fitness has limited support (55). While we found that there were no statistical differences between the growth rate of pseudolysogens and wild type host cells, we found that pseudolysogeny provided surface-level protection against further infection. Furthermore, we found that loss of episomally maintained viruses renders pseudolysogens sensitive to further infection. Hence, viral acquisition comes with a cost (the risk of lytic infection) but is key to enabling surface-level resistance and population level recovery. More work is needed to understand when pseudolysogens become resistant and the cellular mechanisms underlying transient protection to infection. We also found that *Sal. ruber* strains that were sensitive to the EM1 virus were killed in the presence of pseudolysogens. Similar dynamics have been observed with temperate viruses of *Roseobacter*, where lysogens induce and release virions to kill competing sensitive strains in head-to-head experiments (45). We hypothesize that *Sal. ruber* M1 pseudolysogens may switch from a maintenance to an actively infected state, e.g., when episomal numbers are low and/or when resources become available. Again, future work will be needed to elucidate the cellular drivers of lytic initiation and the extent to which intrinsic or extrinsic heterogeneity plays a role in the lysis of pseudolysogens that release EM1 viruses that may then infect and kill competing sensitive strains and potentially initiate new pseudolysogens.

In closing, this study reveals how feedback mechanisms at the cellular scale can transform population dynamics leading to non-monotonic relationships between virus infection pressure and bacterial impacts. Put simply: for strictly lytic viruses, more infection does not always result in more killing. Within the *Sal. ruber* M1-EM1 bacteria-virus model system, we have identified feedback mechanisms that limit lysis, whether because of multiple infections, resource limitation, or the establishment of an episomally maintained virus (i.e., a pseudolysogen). Each of these mechanisms stabilizes virus-microbe population dynamics. We recognize that it is challenging to move from experimental settings to extreme field conditions, hence we are forced - for now – to speculate that pseudolysogeny may be an evolutionarily adaptive strategy for obligately lytic viruses to preserve hosts until environmental conditions improve or host density increases (23, 24, 56). Depending on the durability of viruses extracellularly, such a balance between infection and production may help to explain virus:host ratios ranging from 10 to as high as 300 in the face of fluctuating conditions (14). More broadly, it may be of some value to explore other, related cellular mechanisms that stabilize virus-host population dynamics, enabling reproducible recovery dynamics seen in ecosystems outside hypersaline environments, such as the bloom-bust dynamics observed in oceanic *Emiliania huxleyi* populations after infection by EHV at high MOI (46, 57) or the long-term persistence of crAss-like phage crAss001 and its host (58). Overall, the results from this work highlight the ecological relevance of exploring a continuum of viral infection strategies beyond the dichotomous options of lysis or lysogeny.

## METHODS

### Growth conditions and DNA extraction

*Sal. ruber* strain M1 was grown aerobically with gentle shaking (60 rpm) at 37 °C in 25% SW (sea water) with 0.2% yeast extract (59). *Sal. ruber* M1 was infected with the lytic virus EM1 (27), using a multiplicity of infection (MOI) of 0.01 plaque forming units (PFUs) per cell at time 0 h. Briefly, viruses were mixed with cells at exponential phase (optical density, OD, at 600 nm=0.3) and incubated during 30 minutes at room temperature without shaking to facilitate the adsorption. After the adsorption, the abovementioned culture medium was added up to 25 ml and incubated as previously described. Infections were monitored by OD at 600 nm. Non-infected *Sal. ruber* M1 was grown as the control. All experiments were conducted in triplicate. Another curve was made with the same methodology but using a MOI of 0.1 and 10 replicates in both control and infection.

Bacterial pellets of the initial (C_0_, before virus addition) and final cells (C_F_) of the curves performed using a MOI of 0.01 were obtained by centrifugation at 17.000 xg during 10 min. The cells were washed three times with sterile medium, the supernatant removed and the cell pellets stored at −80 °C. C_0_ and C_F_ DNAs were extracted with the DNeasy Blood & Tissue Kit (Qiagen, Hilden, Germany) following the manufacturer’s protocol, and nucleic acids eluted in 70 μl of milli-Q water and quantified using Qubit 2.0 Fluorometer (Life Technologies, Carlsbad, USA).

### DNA sequencing and bioinformatic analysis

DNA was sequenced on an Illumina HiSeq (2 x 150 bp) at Novogene (Novogene, Seur, UK). Primers and adapters were removed from sequences, and reads were filtered based on quality scores using Trimmomatic v0.36.0 (60). Trimmed reads were assembled using SPAdes v3.13.1 with the trusted option using the reference genome of *Sal. ruber* M1 as scaffold (61). The mean sequencing depth was calculated by a BLASTN-recruitment analysis where the assembled contigs were used as reference.

For the search for hybrid host-virus reads, a BLASTN of the trimmed reads was performed, using the genome of *Sal. suber* M1 of NCBI as a database (GenBank accession number: NZ_CP030364.1). The reads with >95% ID were filtered by best hit and extracted. No horizontal coverage filter was used. Another BLASTN analysis was performed with these reads using the genome of the EM1 virus of NCBI as a database (GenBank accession number: NC_042348.1) with the same parameters. No hybrid host-virus reads were found.

Mutations generated during the infective process were identified comparing reads from C_F_ with the reference contigs assembled from C_0_ using Geneious software 6.1.8 (Biomatters, Auckland, New Zealand) with a minimum variant frequency of 0.01% and a coverage >25% to the mean coverage.

### PCR and epifluorescence microscopy

Cells from C_0_ and C_F_ were diluted and plated on SW 25% with 0.2% yeast extract agar plates (concentration 2%) and incubated at 37 °C for 30-45 days until colonies visualization. Twenty colonies from every point (C_0_ and C_F_) were grown in 2 ml of culture medium at 37 °C and 60 rpm. Once the cultures grew to late exponential phase, 200 µl of each were washed three times by centrifugation (see above). DNA was extracted from washed pellets by boiling at 100 °C for 10 min after resuspending the cells in 80 µl of milli-Q water. Two parallel PCR reactions were performed with each DNA, using specific primers for *Sal. ruber* M1 and the virus EM1 (Supplementary Table 4). PCR reactions were carried out in a final volume of 25 μl, containing: 0.75 μl of 1.5 mM MgCl_2_, 2.5 μl of 10X reaction buffer, 0.5 μl of 10 mM dNTPs, 0.1 μl of Taq polymerase (5 U/μl, Invitrogen, Waltham, USA), 0.5 μl of 10 μM primers, 1 μl of extracted DNA and milli-Q water to complete the final volume. PCR conditions are described in Supplementary Table 5. PCR products were electrophoresed and UV-visualized after staining with ethidium bromide (100 μg/ml).

Three random cultures from C_F_ colonies that were PCR positive for both *Sal*. *ruber* M1 and the EM1 virus (named 1R, 2R and 3R) were selected for the visualization of extracellular virions by SYBR^TM^Gold staining (62). Briefly, 100 µl of the culture were fixed with formaldehyde (4% final concentration) for 1 h at 4 °C. Fixation was stopped by adding phosphate-buffered saline (PBS) 1X up to 1 ml. Twenty microliters (equivalent to 2 µl of sample) were filtered through 0.02 μm Anodisc 25 filters (Whatman Int. Ltd, Maidstone, UK) to retain cells and viruses. Filters were stained with SYBR^TM^Gold (25X) for 15 min, washed twice for 1 min with milli-Q water and visualized with an epifluorescence microscope (Leica, type DM4000B).

### Pulsed-field gel electrophoresis (PFGE) and Southern Blot

One ml of liquid cultures of C_0_, PCR positive for M1, and C_F_, PCR positive for M1 and EM1, were centrifuged and the cell pellets mixed with agarose at 0.8% (final concentration) to obtain plugs, as detailed in Santos et al., 2007 (63). Plugs were incubated overnight at 50 °C in ESP (0.5 M EDTA, pH 9.0, 1% N-laurylsarcosine, 1 mg ml^−1^ proteinase K) for cell lysis and virion disruption and stored at 4 °C. DNA was separated by PFGE on a 1% LE agarose gel in 0.5X TBE buffer, using a Bio-Rad CHEF DR-III System (Bio-Rad, Richmond, USA), under the following conditions: 8–12 s pulse ramp, 6 V/cm, at 14 °C for 30 hrs. After electrophoresis, gels were stained with ethidium bromide (100 μg/ml) for 10 min, washed with distilled water for 30 min, and UV-visualized with a transilluminator.

Agarose gels were transferred to a positively charged nylon membrane (GE Healthcare, Buckinghamshire, UK) as previously described (64). In parallel, DNA from the EM1 virus was labelled to be hybridized against the membrane using the DIG High Prime DNA Labelling and Detection Starter Kit II (Roche, Mannheim, Germany), following manufacturer’s instructions.

### Growth curves, spot tests and adsorption assays

*Sal. ruber* M1 wild type (as a control) and pseudolysogen 1R were grown in liquid, transferred to fresh media (1% v/v) and incubated at 37 °C with gentle shaking at 60 rpm. Cellular density was measured by OD at 600 nm. The growth curves of *Sal. ruber* M1 wild type and pseudolysogen 1R were compared for statistical differences using 99% confidence intervals obtained through the Delta method (*Modeling Supplementary Information: Sal. ruber M1 wild-type and 1R (pseudolysogen) growth comparison*).

To test the resistance/susceptibility of *Sal. ruber* M1 wild type and pseudolysogen 1R to the EM1 virus, both hosts were exposed to the virus by a spot test. Four ml of molten 0.7% top agar of 25% SW with 0.2% yeast extract were mixed with 500 μl of bacterial cultures at exponential phase and plated on solid medium. Once solidified, 3 μl of the EM1 virus, tittered at 10^10^ PFUs/ml and diluted 10^2^, 10^4^ and 10^6^ times, were added to the lawn. The spotted plates were left to dry and incubated at 37 °C for 10 days. The detection of a clearance zone was interpreted as evidence of viral lysis while the areas exhibiting growth were attributed to resistance against EM1.

An adsorption assay was performed with the EM1 virus and *Sal. ruber* M1 wild type. Cultures in exponential phase were diluted with fresh medium to an optical density of 0.3 and 2×10^8^ cells were mixed with 2×10^6^ PFU (MOI=0.01) or 2×10^9^ PFU (MOI=10) of the EM1 virus. This was considered as the initial time. It was incubated at 37 °C and 60 rpm. Aliquots of 150 μl were taken along 2 h (at 15, 30, 45, 60, 90 and 120 min), centrifuged at 17.000 xg for 8 min, and 120 μl of the cells-free supernatant were stored in ice. The number of infective free viruses was measured by plaque assay, diluting the supernatants in sterile SW 25% and mixing 100 μl of each with 500 μl of *Sal. ruber* M1 wild type in exponential phase. Four ml of top agar were added and this was plated on solid plaques. After solidifying, the plates were incubated at 37 °C for 10 days until plaques were visible. The number of free infective viruses (PFU/ml) were counted and represented. A control of the viral decay of the EM1 virus was added, in which the virus was incubated without host. All the experiments were conducted in triplicate.

To get more information about the resistance mechanism, an adsorption assay was performed with the EM1 virus and *Sal. ruber* M1 wild type and pseudolysogen 1R with a MOI of 0.01 as described above.

To calculate the adsorption rate, the graph of the free infective viruses (PFU/mL) was plotted with respect to time and fit to an exponential function. As the assay was conducted within a 120 min time period and the latent period of the EM1 virus is around 21-22 h, we can assume that there is negligible killing and the susceptible host population stays constant at *S* = 2×10^8^ cells/ml. We fit the adsorption data to the function *V* = *V*_0_ *e*^−ɸ*St*^ and get an adsorption rate ɸ ≈ 4×10^-10^ ml/h (Fig. S11).

### Infection curves at varying MOIs and revival of lysis in pseudolysogens

*Sal. ruber* M1 wild type was grown at 37 °C and 60 rpm. The EM1 virus was added at 67 h at middle exponential phase. Cultures were infected with a MOI of 0.01, 0.1, 1, 5 and 10 plaque forming units (PFUs) per cell in separate experiments. Infections were monitored by OD at 600 nm. *Sal. ruber* M1 was grown without viruses as a control. All experiments were conducted in triplicate.

A total of 2×10^8^ cells from the end of each infection curve were transferred to 4 ml of fresh medium (SW 25% with 0.2% yeast extract). This caused all cultures to be diluted to an OD of approximately 0.1. Cultures were monitored by OD at 600 nm.

### Numerical Simulations

All the numerical simulations were performed on Python 3.11.5 through the Jupyter IDE. To solve the model ODEs and obtain the simulated results, we used the numerical integration module *odeint* from the *SciPy* package. To convert the experimental OD measurements to concentrations, we fit a calibration curve using the *curve_fit* module from the *SciPy* package. All the code that can reproduce the simulated results seen in this paper are available on the Github: https://github.com/aranilah/SalruberM1_EM1_recovery/tree/main

### Transfers of pseudolysogen 1R to fresh media, PCR and qPCR

In order to assess if the EM1 virus was with generations, pseudolysogen 1R was transferred four times to fresh media. Briefly, pseudolysogen 1R was transferred in triplicates to fresh media (1 % v/v) and incubated at 37 °C with shaking at 60 rpm, measuring the growth by OD at 600 nm. At the late exponential phase cultures were transferred again to fresh media until a total of 4 transfers. A spot test of each transfer was also carried out as previously mentioned to check if the different transfers of pseudolysogen 1R were still resistant to the virus. Moreover, 100 μl of each transfer were fixed with formaldehyde (0.5% final concentration to avoid PCR inhibition) for 1 h at 4 °C. Fixation was stopped by adding phosphate-buffered saline (PBS) 1X up to 1 ml.

To eliminate free viruses, assuming that viruses could have been spontaneously released during the transfer periods, the fixed samples were centrifuged at 17.000 xg during 10 min and washed with PBS five times. PCR reactions for *Sal. ruber* M1 and the EM1 virus were carried out as mentioned above, with 1 μl of the fixed and washed cells as DNA template (first, we checked that the PCR initial thermal shock was sufficient for cell lysis; data not shown).

Once the presence of EM1 was detected in all the transfers, a qPCR was performed to monitor virus concentration along the subsequent transfers. The qPCR assay was conducted using TaqMan hydrolysis probes (Supplementary Table 4) labeled with hexachlorofluorescein (for *Sal. ruber* M1) and fluorescein (for EM1). The experiment was carried out using the standard run in a StepOnePlus™ PCR System (Life Technologies, Carlsbad, USA) in a 10 μl reaction mixture with PrimeTime™ Gene Expression Master Mix (Integrated DNA Technologies, Coralville, USA). The reaction contained: 5 μl of 2X Master Mix, 0.2 μl of each 10 μM primer, 0.2 μl of 10 μM TaqMan probe, 1 μl of fixed sample and milli-Q water to complete volume. Conditions are detailed in Supplementary Table 6. The results were analyzed with the Applied Biosystems StepOne™ Instrument program. All samples were run in triplicate (including the standards and negative controls).

### Plating of pseudolysogen 1R, mutation screening and spot test

Pseudolysogen 1R at late exponential phase was plated on solid medium (SW 25% with 0.2% yeast extract and 2% agar) to obtain individual colonies. After two months of incubation, 50 colonies developed, which were picked and transferred to 1 ml of liquid media each. The presence of the EM1 virus and the resistance/susceptibility was checked by PCR and spot test as previously described.

To check if the colonies had the mutation affecting the sodium/glucose cotransporter, primers were designed to amplify the specific region (Supplementary Table 7). DNAs of the 6 selected colonies picked in 1 ml of medium and *Sal. ruber* M1 wild type (as a control of the mutation) were extracted by boiling as previously described and PCR reactions done as described previously. PCR conditions are described in Supplementary Table 8. PCR products were visualized under ultraviolet light after staining with ethidium bromide (100 μg/ml). Products were sequenced with Sanger method at Stab Vida (Stab Vida, Caparica, Portugal) and analyzed with Geneious software 6.1.8 (Biomatters, Auckland, New Zealand).

Eight strains of *Sal. ruber* (M1, M8, M31, P13, P18, SP38, SP73 and RM158) were exposed to pseudolysogen 1R through spot test as previously described, adding spots of 3 μl of 1R at late exponential phase and diluted 10^2^, 10^4^ and 10^6^ times on the double layer formed by the strains.

### One step growth curve

Burst size was determined after carrying out a one-step growth experiment as detailed in Kropinski, 2018 (65) (Supplementary Table 9).

## Supporting information

Mutation Supplementary Information

Modeling Supplementary Information

## ACKNOWLEDGEMENTS

We warmly thank to Julia Esclapez and Laura Matarredona for their help with the Southern Blot. We thank Cristina López for providing the initial infection curves. This research was supported by the projects “Virhost” CIPROM/2021/006 (PROMETEO 2022, Generalitat Valenciana) and METACIRCLE PID2021-126114NB-C41 (Spanish Ministry of Science and Innovation). R.S.M. received funding for his doctoral thesis from the Spanish Ministry of Science and Innovation PRE2019-087998. R.S.M., F.S. and J.A. are members of the National Excellence Network FAGOMA (RED2022-134837-T). This project was supported, in part, by Simons Foundation Grant 722153 (to J.S.W.) and by the Chaires Blaise Pascal program of the Île-de-France Region (to J.S.W.).

## AUTHOR CONTRIBUTIONS

R.S.M., J.S.W., F.S. and J.A. conceived and designed the study. R.S.M. performed the experiments and analyzed the sequences under the inputs and supervision of M.K., F.S., and J.A.. A.A. and J.S.W. designed and implemented the mathematical model. A.A. wrote the code and performed the simulations and statistical analysis. R.S.M. and A.A. drafted the original manuscript. All authors contributed to manuscript final writing and approved the final paper.

## DATA AVAILABILITY

The raw files used for mutation analysis were deposited in the NCBI database with BioProject accession ID PRJNA1136859. Raw sequences of the Sanger sequencing of the mutation affecting the sodium/glucose cotransporter can be found in the *Mutations Supplementary Information*

## CODE AVAILABILITY

The code used in the study is available via GitHub at: https://github.com/aranilah/SalruberM1_EM1_recovery/tree/main

## ETHICS DECLARATIONS

The authors declare no competing interests.

## EXTENDED DATA

**Extended Data Fig. 1.**
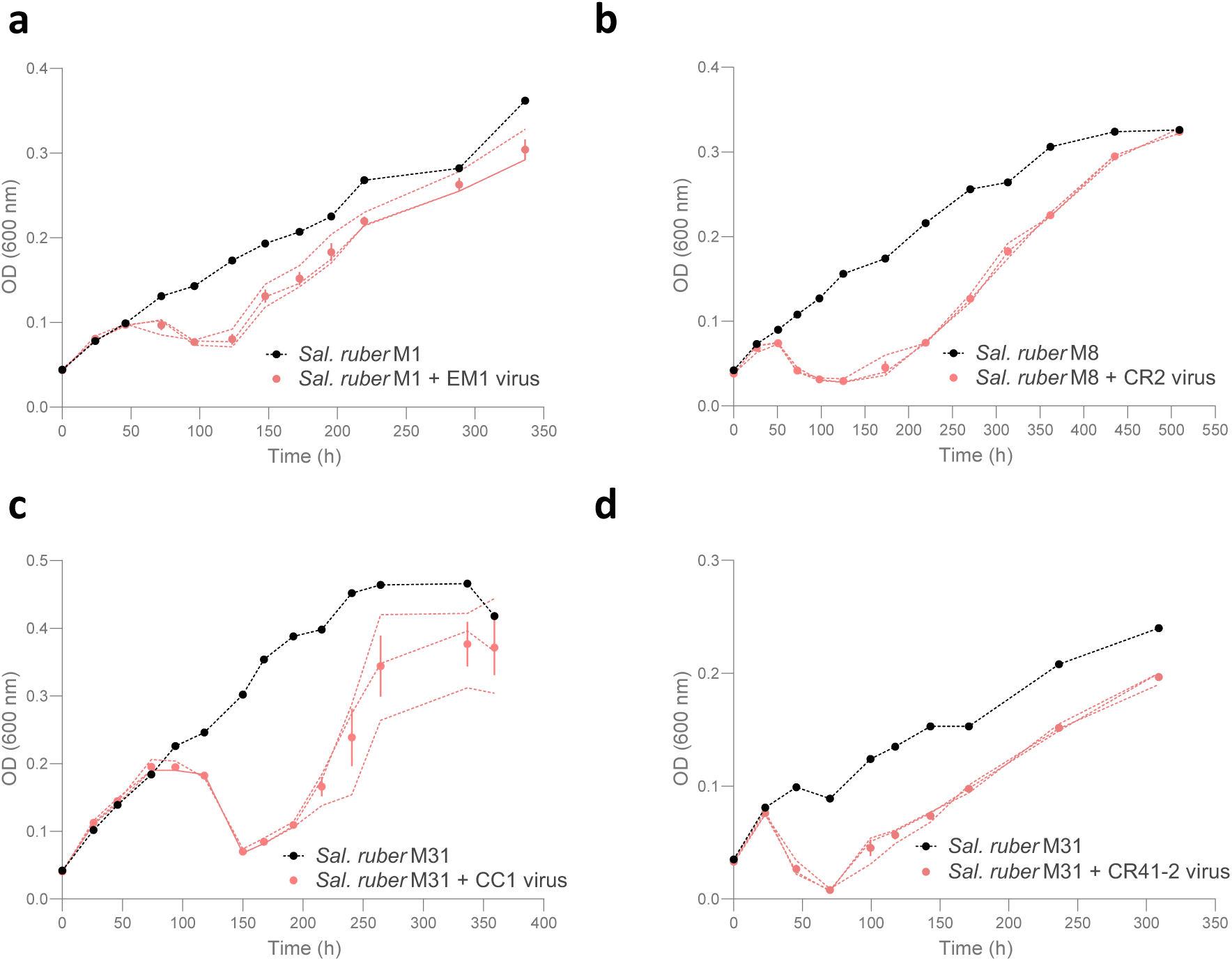
Infection curves of different *Sal. ruber* strains and its viruses. The black dotted line represents the control without viruses and the pink dotted lines represent each replicate of the infected culture. In the infected cultures, the mean is represented by a pink dot and the bars represent the standard error. These curves were made with different strains of *Sal. ruber* and different viruses (**a**, strain M1 and EM1 virus; **b**, strain M8 and CR2 virus; **c**, strain M31 and CC1 virus; **d**, strain M1 and CR41-2 virus). The control has only one replicate, while the infection has 3 replicates.

**Extended Data Fig. 2.**
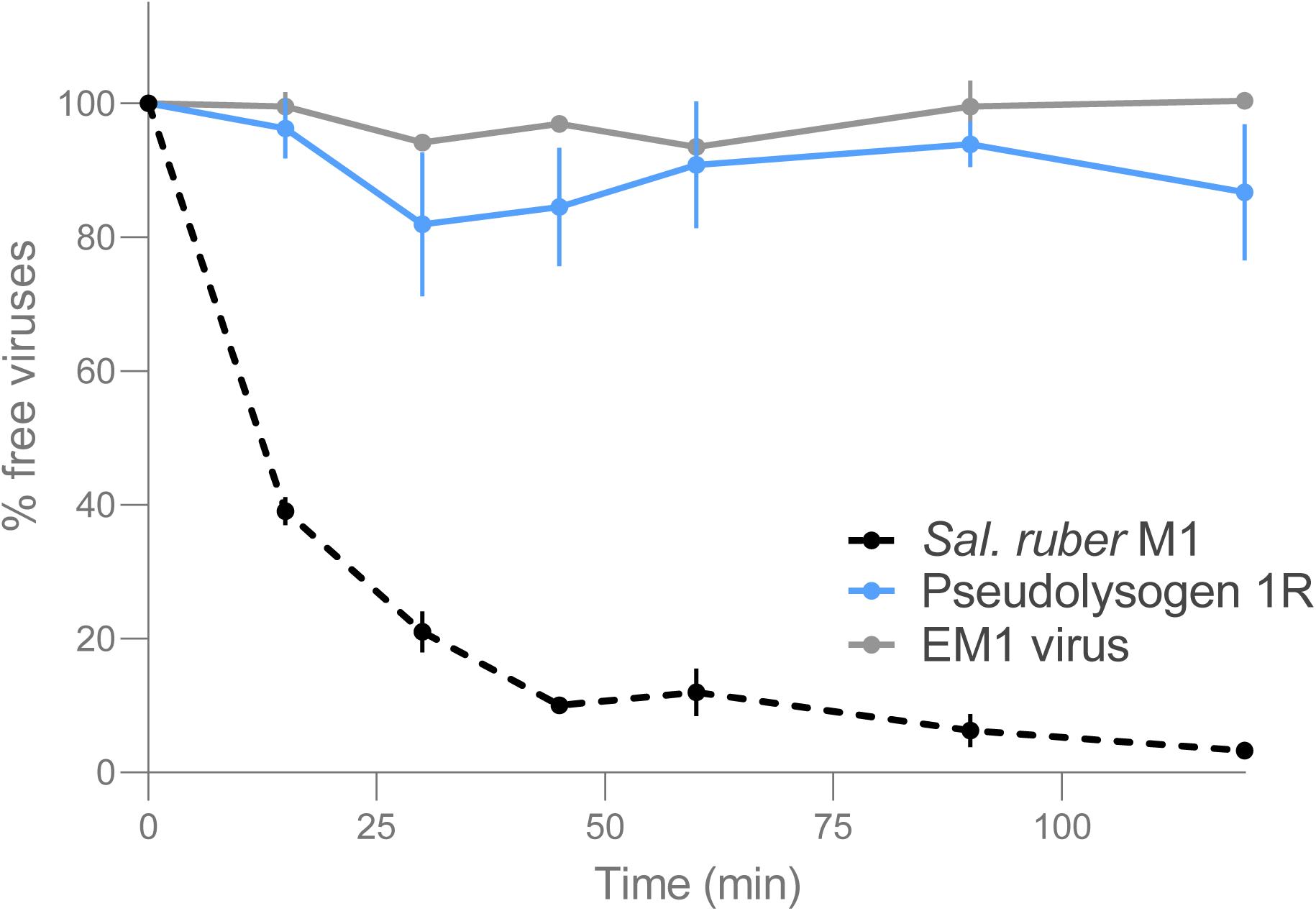
Pseudolysogens develop resistance to EM1 at external surface level. An adsorption assay of the EM1 virus to *Sal. ruber* M1 wild type (black dotted line) and pseudolysogen 1R (blue line) was carried out with a MOI of 0.01, quantifying the quantity of extracellular viruses through PFU/ml during 120 minutes. The experiment was also carried out with the virus alone as a decay control (grey line). The experiment was performed by triplicate. Error bars represent the standard error.

**Extended Data Fig. 3.**
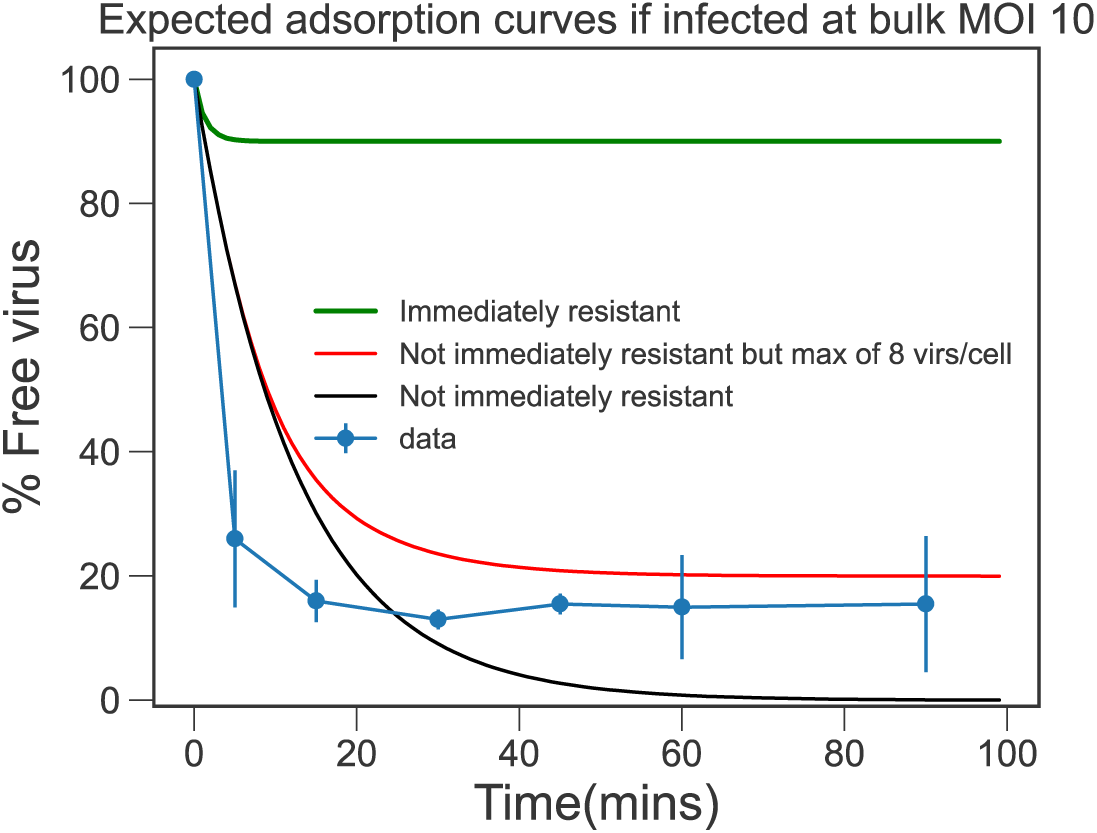
Expected adsorption results with superinfection. The simulated adsorption dynamics when only a single virus can infect a cell (*k=1, green)*, when eight viruses at maximum can infect a cell *k= 8* viruses (*k=8, red*), and when viruses can continuously infect a cell with no limit (*black*). The dynamics when *k=8* seem to match the experimental adsorption dynamics observed in Figure 3d. In reality, the reasoning behind the limit to the number of viruses per cell could be due to competition for receptor sites, or perhaps explicit spatial limitations.

**Extended Data Fig. 4.**
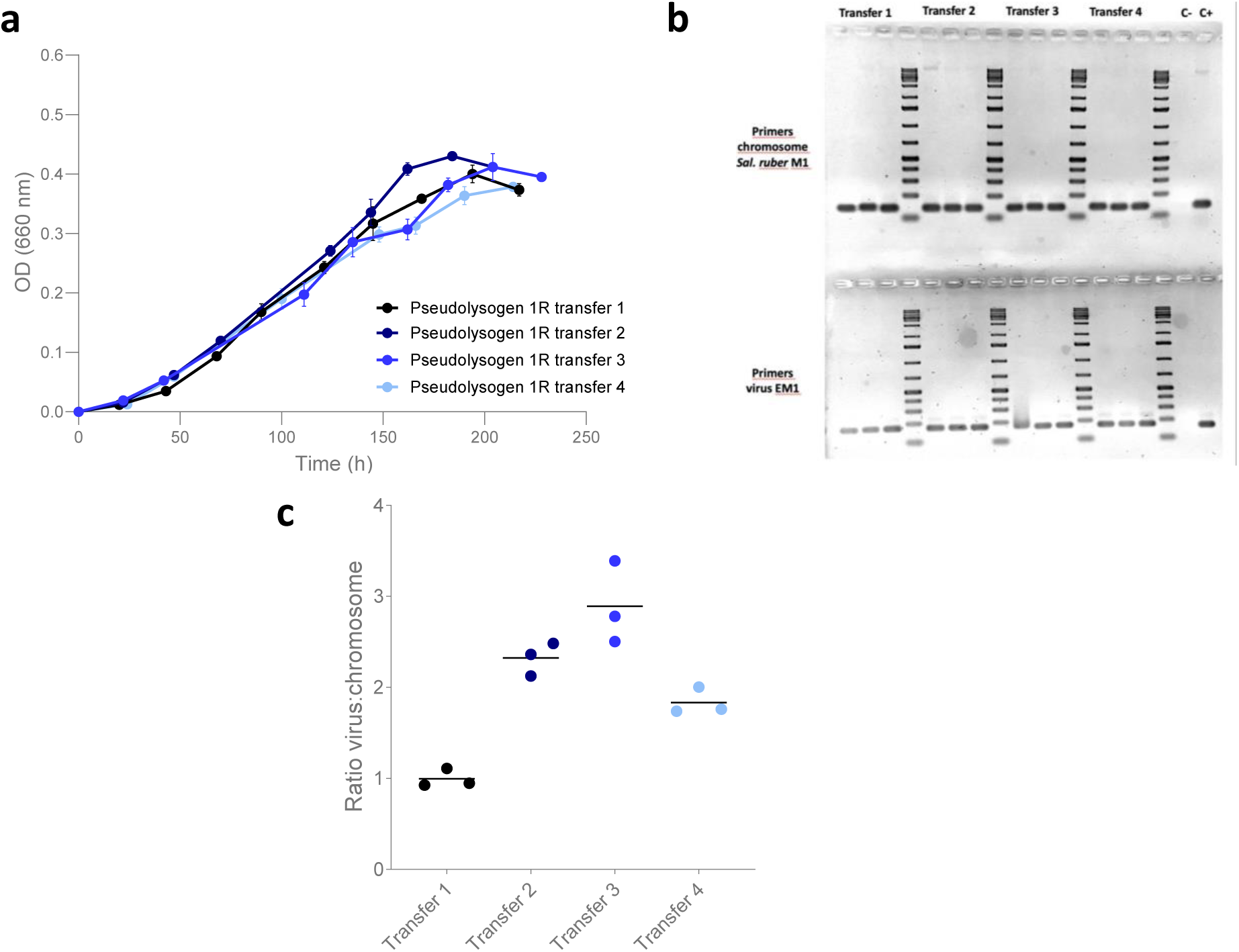
The EM1 virus is maintained at the population level across time. **a**, Growth curves of the different transfers of pseduolysogen 1R. The points taken for new transfers, spot tests, PCR and qPCR are marked with a grey oval. It was transferred by triplicate. Error bars represent the standard error of the replicates. **b**, PCR of the three replicates of the transfers of pseudolysogen 1R shown above. The PCR was performed with primers for *Sal. ruber* M1 (upper part) and the EM1 virus (bottom part). **c**, qPCRs were performed on each of the 4 transfers to quantify the ratio of EM1 virus genome/chromosome of *Sal. ruber* M1 in the culture. Biological replicates are represented with dots. The mean is represented as a horizontal line.

**Extended Data Fig. 5.**
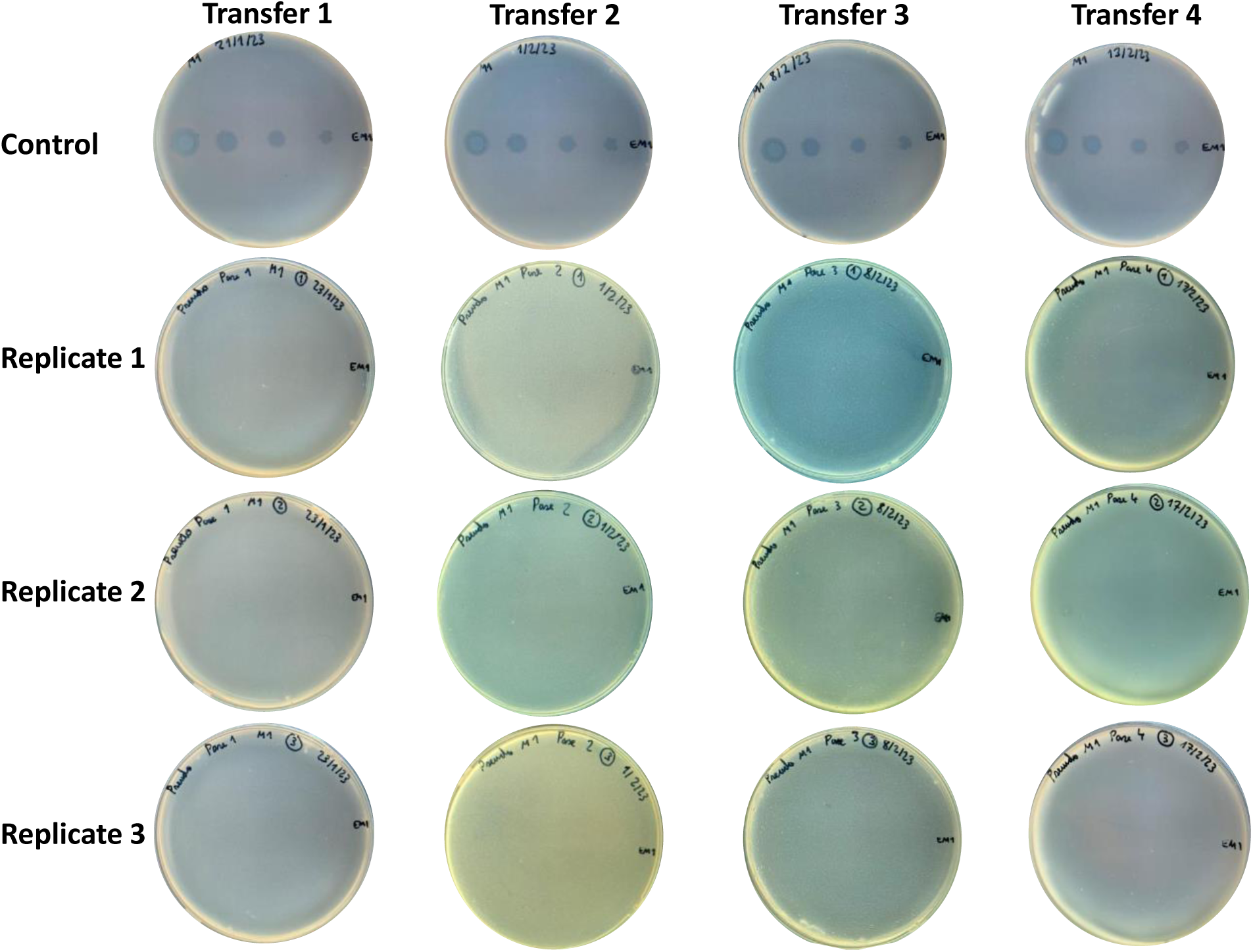
Resistance to the EM1 virus is also maintained over time in the pseudolysogens. Spot tests of the three replicates of the transfers of pseudolysogen 1R to assess the susceptibility to the EM1 virus. All the transfers of pseudolysogen 1R were plated in solid medium and exposed to spots of the EM1 virus tittered at 10^10^ PFUs/ml and diluted 10^2^, 10^4^ and 10^6^ times. A control of *Sal. ruber* M1 wild type (upper part) was included in each transfer to test the infectivity of the virus.

**Extended Data Fig. 6.**
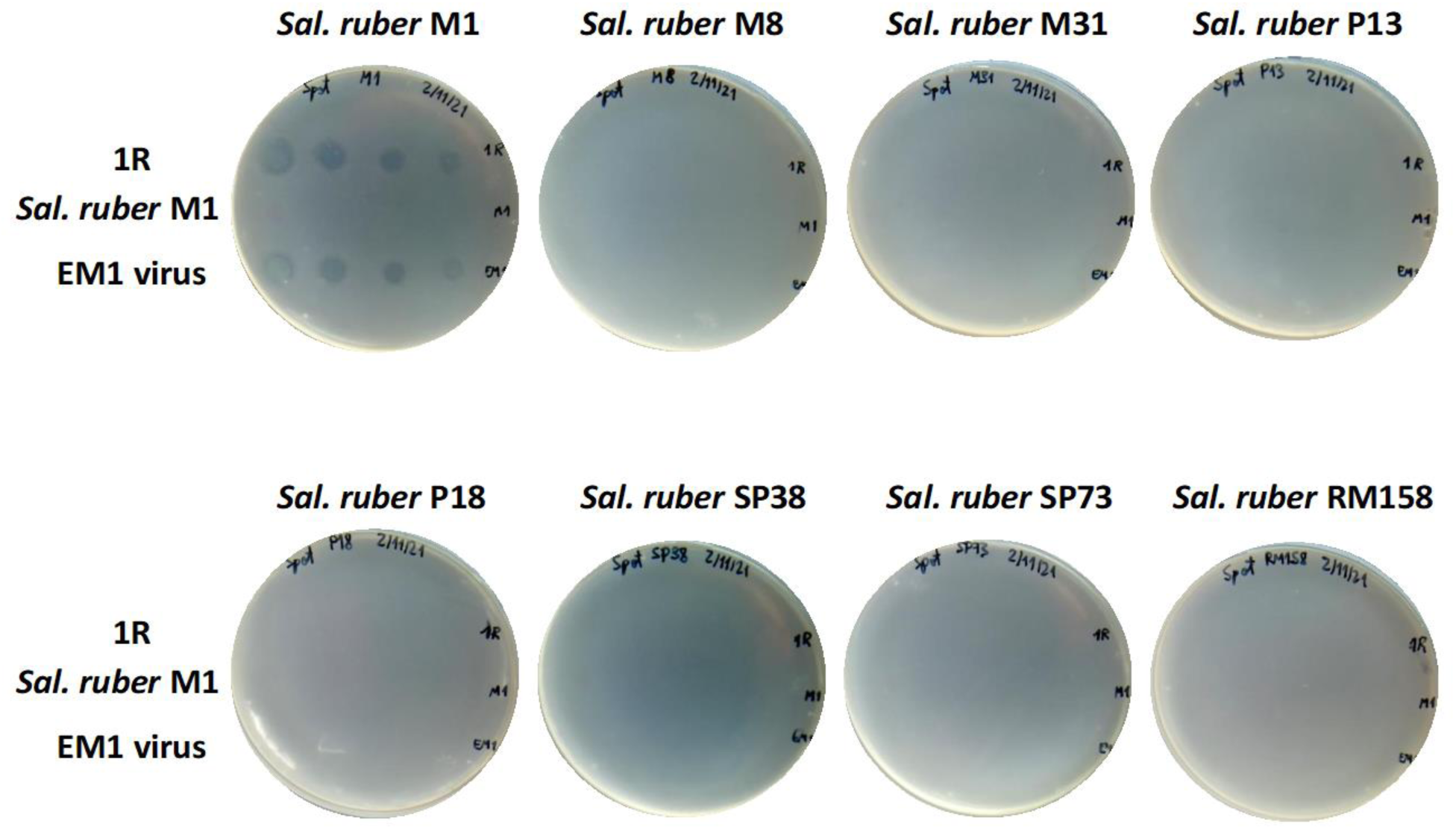
Pseudolysogens kill other virus-susceptible strains. Spot test of eight wild type strains of *Sal. ruber* against the pseudolysogen 1R. All the cultures were plated in solid medium and exposed to spots of pseudolysogen 1R diluted 10^0^, 10^2^, 10^4^ and 10^6^ times. Spots of *Sal. ruber* M1 wild type and the EM1 virus were also added as a control.

