## Supplementary material for "Episomal virus maintenance enables bacterial population recovery from infection and virus-bacterial coexistence": Mutation Supplementary Information

**Supplementary Table 1**

| Product | Position | Protein Effect |
|---|---|---|
| ATP-dependent helicase/deoxyribonuclease subunitB | 56473 | Substitution |
| Hypothetical protein | 858339 | None |
| Sodium/glucose cotransporter | 1784378 | Extension |
| Non-codificant region | 227,834 | None |
| Orotate phosphoribosyltransferase | 2371568 | Substitution |
| Hypoxanthine-guanine phosphoribosyltransferase | 3465140 | None |

**Supplementary Table 2**

| gene_ID | gene_start | gene_end | gene_length | Protein prediction |
|---|---|---|---|---|
| gene_1 | 1 | 237 | 237 | Hypothetical protein |
| gene_2 | 729 | 914 | 186 | DNA-binding HTH domain-containing protein |
| gene_3 | 911 | 2371 | 1461 | Hypothetical protein |
| gene_4 | 2397 | 3746 | 1350 | Structural protein |
| gene_5 | 3829 | 5040 | 1212 | Hypothetical protein |
| gene_6 | 5045 | 6187 | 1143 | Putative head morphogenesis protein |
| gene_7 | 6396 | 7133 | 738 | Head scaffolding protein |
| gene_8 | 7209 | 8033 | 825 | Major head protein |
| gene_9 | 8061 | 8498 | 438 | Hypothetical protein |
| gene_10 | 8502 | 8831 | 330 | Minor capsid protein |
| gene_11 | 8835 | 9149 | 315 | Neck protein Ne1 |
| gene_12 | 9151 | 9579 | 429 | Phage minor tail protein |
| gene_13 | 9579 | 10418 | 840 | Structural protein |
| gene_14 | 10498 | 10851 | 354 | Hypothetical protein |
| gene_15 | 10941 | 11210 | 270 | Hypothetical protein |
| gene_16 | 11200 | 13431 | 2232 | Minor tail protein |
| gene_17 | 13450 | 18009 | 4560 | Long tail fiber proximal subunit |
| gene_18 | 18299 | 18661 | 363 | Peptidase |
| gene_19 | 18658 | 19095 | 438 | Hypothetical protein |
| gene_20 | 19076 | 19663 | 588 | Hypothetical protein |
| gene_21 | 19677 | 20018 | 342 | Transcriptional repressor |
| gene_22 | 20679 | 21395 | 717 | DNA-binding HTH domain-containing protein |
| gene_23 | 21482 | 21718 | 237 | Hypothetical protein |
| gene_24 | 21822 | 22013 | 192 | Hypothetical protein |
| gene_25 | 22267 | 22530 | 264 | Hypothetical protein |
| gene_26 | 22542 | 23498 | 957 | Nuclease domain-containing protein |
| gene_27 | 23905 | 24336 | 432 | Structural protein |
| gene_28 | 24409 | 24861 | 453 | Primosome PriB/single-strand DNA-binding |
| gene_29 | 24893 | 25141 | 249 | Single stranded DNA-binding protein |
| gene_30 | 25316 | 25678 | 363 | DNA-binding HTH domain-containing protein |
| gene_31 | 25828 | 26331 | 504 | Putative nuclease |
| gene_32 | 26495 | 26761 | 267 | Hypothetical protein |
| gene_33 | 26758 | 26970 | 213 | Hypothetical protein |
| gene_34 | 26967 | 27326 | 360 | RusA-like Holliday junction resolvase |
| gene_35 | 27337 | 28434 | 1098 | DNA polymerase III beta subunit |
| gene_36 | 28494 | 30764 | 2271 | Topoisomerase-primase domain-containing protein |
| gene_37 | 30955 | 31755 | 801 | GIY-YIG nuclease family protein |
| gene_38 | 31978 | 33180 | 1203 | DNA modification methylase |
| gene_39 | 33534 | 34505 | 972 | Deoxynucleoside monophosphate kinase |
| gene_40 | 34600 | 34959 | 360 | Hypothetical protein |
| gene_41 | 34968 | 35684 | 717 | Terminase small subunit |

**Supplementary Table 3**

| Sample | Replicate | Origin | Nt | Seq. depth |
|---|---|---|---|---|
| M1 F | 1 | Cromosomal | 276,949 | 112.8 |
| | | Cromosomal | 819835 | 92.9 |
| | | Plasmid pSR116 | 116032 | 55.9 |
| | | Plasmid pSR66 | 66225 | 36.2 |
| | | Plasmid pSR61 | 61538 | 32.0 |
| | | Plasmid pSR10 | 1,261 | 279.0 |
| | 2 | Cromosomal | 276,948 | 104.8 |
| | | Cromosomal | 819835 | 84.9 |
| | | Plasmid pSR116 | 116032 | 54.7 |
| | | Plasmid pSR66 | 66225 | 32.8 |
| | | Plasmid pSR61 | 61538 | 29.9 |
| | | Plasmid pSR10 | 1,261 | 240.8 |
| | 3 | Cromosomal | 276,944 | 117.3 |
| | | Cromosomal | 819835 | 95.5 |
| | | Plasmid pSR116 | 116032 | 53.7 |
| | | Plasmid pSR66 | 66225 | 36.3 |
| | | Plasmid pSR61 | 61538 | 33.3 |
| | | Plasmid pSR10 | 1,261 | 270.0 |
| M1 EM1 F | 1 | Cromosomal | 3528100 | 104.3 |
| | | Plasmid pSR116 | 116032 | 113.1 |
| | | Plasmid pSR66 | 66225 | 85.2 |
| | | Plasmid pSR61 | 61538 | 95.4 |
| | | Plasmid pSR10 | 1,261 | 337.9 |
| | | Virus EM1 | 35574 | 154.1 |
| | 2 | Cromosomal | 3528100 | 121.1 |
| | | Plasmid pSR116 | 116032 | 107.9 |
| | | Plasmid pSR66 | 66225 | 83.9 |
| | | Plasmid pSR61 | 61538 | 93.1 |
| | | Plasmid pSR10 | 1,261 | 355.7 |
| | | Virus EM1 | 35574 | 133.6 |
| | 3 | Cromosomal | 3528104 | 122.1 |
| | | Plasmid pSR116 | 116032 | 139.2 |
| | | Plasmid pSR66 | 66225 | 102.9 |
| | | Plasmid pSR61 | 61538 | 113.8 |
| | | Plasmid pSR10 | 1,261 | 395.1 |
| | | Virus EM1 | 35574 | 151.3 |

### Supplementary Table 4

| *Sal. ruber* strain / Virus | Primer / Probe | Sequence |
|---|---|---|
| *Sal. ruber* M1 | Forward | TTCGGCCTGCCTTACTCTTT |
| | Reverse | TTTACCGTCCCCAACCAAGT |
| | Taqman probe | AGCGGAACTGCAAAGACAAGGACATGAGT |
| EM1 virus | Forward | GGTCGCGGGGCTTAATATC |
| | Reverse | CGTGTTGTTCGTTCCCCTTT |
| | Taqman probe | ACTGCCAACCCCGACGACTCCAC |

### Supplementary Table 5

| Nº of cycles | Temperature | Time |
|---|---|---|
| 1 cycle | 95°C | 2 minutes |
| 30 cycles | 95°C | 15 seconds |
| | 59ºC | 15 seconds |
| | 72ºC | 45 seconds |
| 1 cycle | 72ºC | 10 minutes |
| 1 cycle | 4ºC | ∞ |

### Supplementary Table 6

| Nº of cycles | Temperature | Time |
|---|---|---|
| 1 cycle | 95°C | 20 seconds |
| 40 cycles | 95°C | 1 second |
| | 60ºC | 20 seconds |
| 1 cycle | 4ºC | ∞ |

### Supplementary Table 7

| Region | Primer / Probe | Sequence |
|---|---|---|
| Sodium/glucose cotransporter | 205F | GGGGAAGGAGCAGCTTCAG |
| | 414R | AGCTGATGTACCGCAACGAC |

### Supplementary Table 8

| Nº of cycles | Temperature | Time |
|---|---|---|
| 1 cycle | 95°C | 2 minutes |
| 30 cycles | 95°C | 15 seconds |
| | 59ºC | 15 seconds |
| | 72ºC | 1 min 15 seconds |
| 1 cycle | 72ºC | 10 minutes |
| 1 cycle | 4ºC | ∞ |

### Supplementary Table 9

| | PFU/ml | | | | |
|---|---|---|---|---|---|
| Time (h) | Replicate 1 | Replicate 2 | Replicate 3 | Mean | SD |
| 0 | 140 | 150 | 140 | 1.43E+02 | 5.77E+00 |
| 17 | 180 | 200 | 150 | 1.77E+02 | 2.52E+01 |
| 19 | 250 | 270 | 230 | 2.50E+02 | 2.00E+01 |
| 21 | 280 | 250 | 290 | 2.73E+02 | 2.08E+01 |
| 23 | 450 | 780 | 620 | 6.17E+02 | 1.65E+02 |
| 25 | 1050 | 1160 | 1000 | 1.07E+03 | 8.19E+01 |
| 27 | 1860 | 1950 | 1720 | 1.84E+03 | 1.16E+02 |
| 29 | 1940 | 1900 | 2010 | 1.95E+03 | 5.57E+01 |
| 31 | 1990 | 2140 | 1940 | 2.02E+03 | 1.04E+02 |

| Free PFU/ml after initial adsorption | | | | |
|---|---|---|---|---|
| Replicate 1 | Replicate 2 | Replicate 3 | Mean | SD |
| 40 | 30 | 20 | 30 | 10 |

| Burst size |
|---|
| 11 |
