## Supplementary tables for "Episomal virus maintenance enables bacterial population recovery from infection and virus-bacterial coexistence"

To get information about the mutations produced during the infection, all the replicates of $C_F$ were compared with $C_0$. Mutations with a frequency above 0.01% as our threshold were included.

There were notable differences in the number of variations generated between infected and non-infected cultures in the chromosome, with a higher number of mutations in infected cultures (79252 variations as average in infected cultures and 74465 in non-infected cultures). This could be attributed to the selective pressure generated by the virus. In both cases these mutations tend to accumulate in specific areas of the genome (Supplementary Fig. 1).

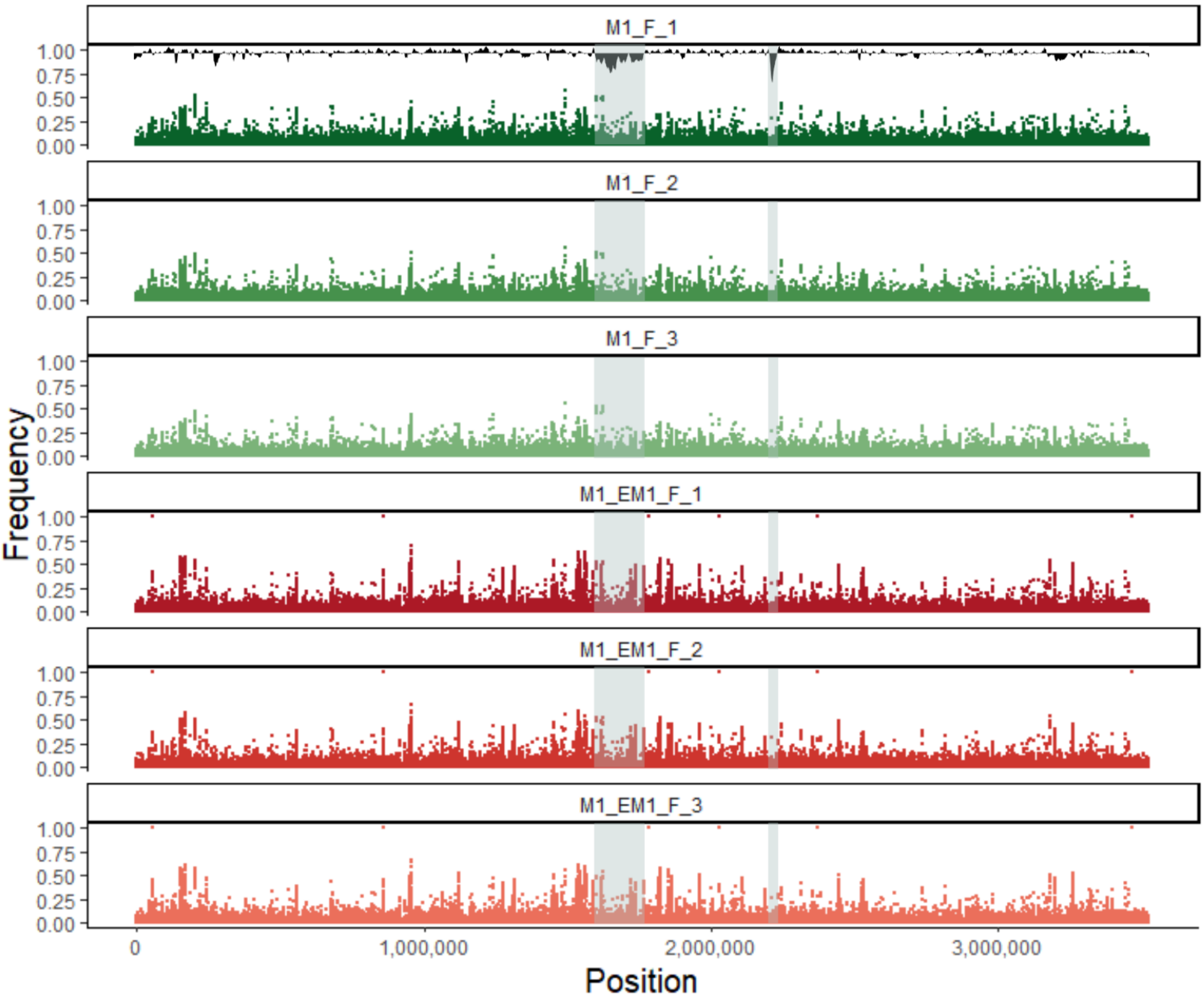


(legend in the next page)

Supplementary Fig. 1. Mutations in the chromosome of *Sal. ruber* M1 after a viral infection are heterogeneously distributed. Distribution of the mutations (dots) along the chromosome after the infection curve. The x-axis represents the length of the chromosome and the y-axis the frequency of each mutation. The first three graphs (top, green dots) refer to the 3 replicates of final cells of the non-infected culture and the last three (bottom, red dots) to the 3 replicates of final cells of the infected culture. In the first graph (top) is represented the %GC content, normalized to the genome mean %GC. Hypervariable zones of *Sal. ruber* M1, that correspond to areas of low %GC, are marked in grey

Only 6 mutations showed a frequency close to 100% (Supplementary Table 1). Of these 6 mutations, 3 supposed a non-synonymous change, affecting an ATP-dependent helicase/deoxyribonuclease, an orotate phosphoribosyltransferase and a sodium/glucose cotransporter. The first two are related to helicase/exonuclease activities and to the synthesis of pyrimidines, respectively, so they do not seem to have a direct relationship with virus resistance. Nevertheless, the mutation affecting the stop codon in the synthesis of the sodium/glucose cotransporter can be involved in the resistance, as it affects a transmembrane protein that could be one of the virus receptors. This mutation was further examined later, showing that it was unrelated to resistance.

Parallelly, to check whether the resistance was due only to the virus, pseudolysogen 1R was plated on solid media, assuming that part of this population would not have the virus. Just a few tiny colonies were obtained, of which only 17 could be picked and transferred to 1 ml of liquid media. The presence of the virus was checked by PCR, showing that 13 of the 17 colonies retained the virus (Supplementary Fig. 2a). The resistance-susceptibility of each picked colony was tested by spot test, which revealed that those cells that did not have the virus were sensitive (Supplementary Fig. 2b), verifying that the resistance was due to the virus acquisition.

Coming back to the mutation affecting the stop codon in the synthesis of the sodium/glucose cotransporter (see above), this region was amplified by PCR and sequencing in the 16 colonies (one of the colonies presented a sequencing error) and the M1 wild type. This showed that both resistant virus carriers and virus-free susceptible colonies had the mutation, confirming that it was not involved in the resistance (Supplementary Fig. 2c).

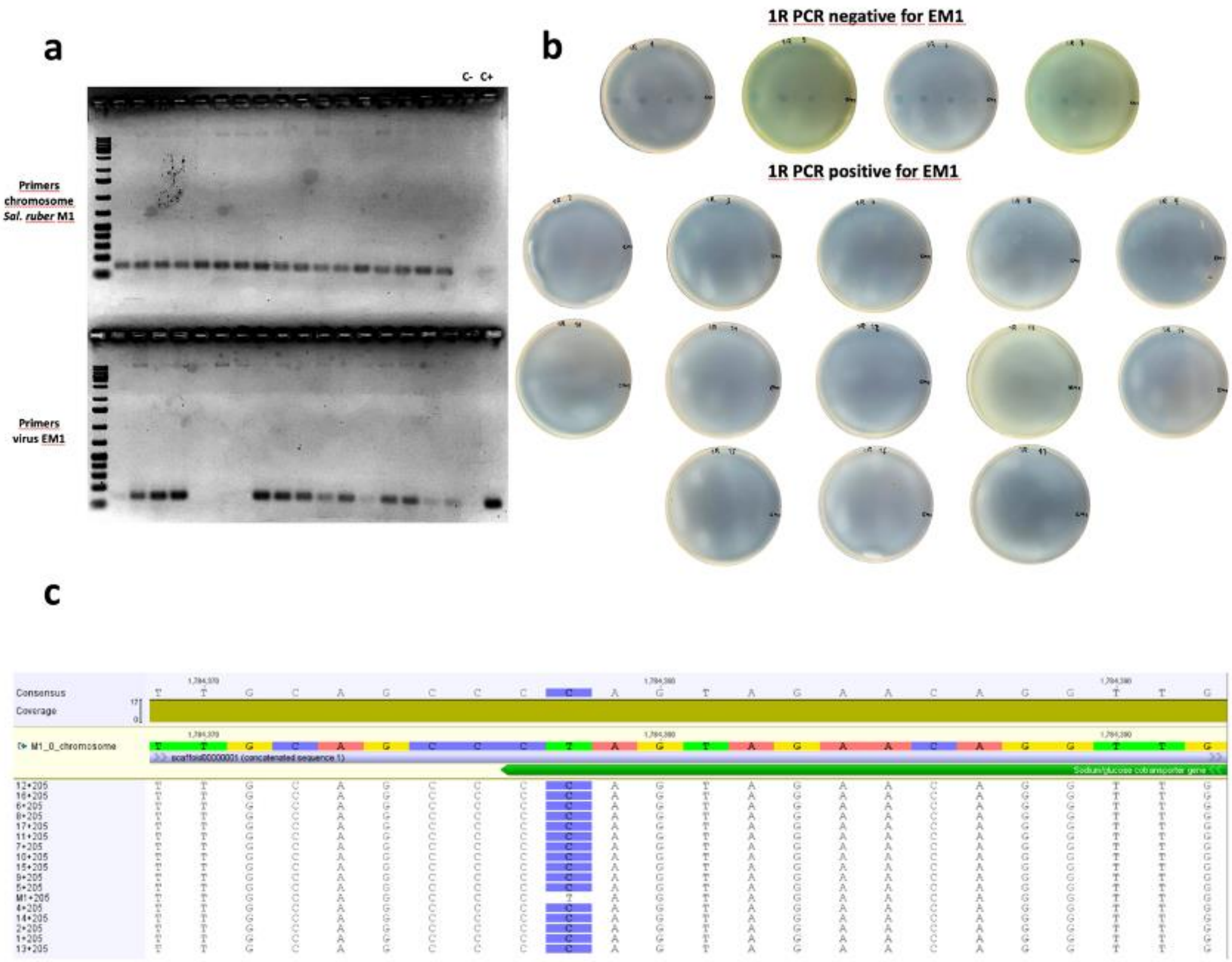


Supplementary Fig. 2. The loss of the virus implies a return to susceptibility**. a,** PCR with the primers of *Sal ruber* M1 (upper part) and EM1 virus (bottom part) from the 17 colonies coming from 1R. **b,** Spot test of the 17 colonies to test the susceptibility to the virus EM1. All the cultures were plated in solid medium and exposed to spots of the virus EM1 tittered at $10^{10}$ PFUs/ml and diluted $10^2$, $10^4$ and $10^6$ times. **c,** Alignment of the amplicons of the region belonging to the sodium/glucose cotransporter mutation of the 16 colonies that could be sequenced by Sanger against the *Sal. ruber* M1 genome. It can be seen that in all of them there is a thymine instead of a cytosine (marked in purple), except for one of them, which is the wild type control strain.

To gain a more detailed insight into the mutations of the revertant pseudolysogens and to ensure that it was the virus protecting the pseudolysogens, the complete genome of 3 colonies (see above) derived from pseudolysogen 1R that retained the virus (resistant) and 3 that lost it (susceptible) were sequenced.

The number of total mutations generated varied considerably between replicates, with no clear pattern found. As in the first sequencing, there was an accumulation of mutations in specific areas of the genome in both the pseudolysogen that retained the virus the and pseudolysogens that lost the virus (Supplementary Fig. 3).

We focused on examining mutations close to 100% frequency. In all cases, the 6 mutations generated in the previous infectious process were maintained (see above), confirming that these mutations were not involved in virus resistance. no other mutations of relevance shared between replicates were found.

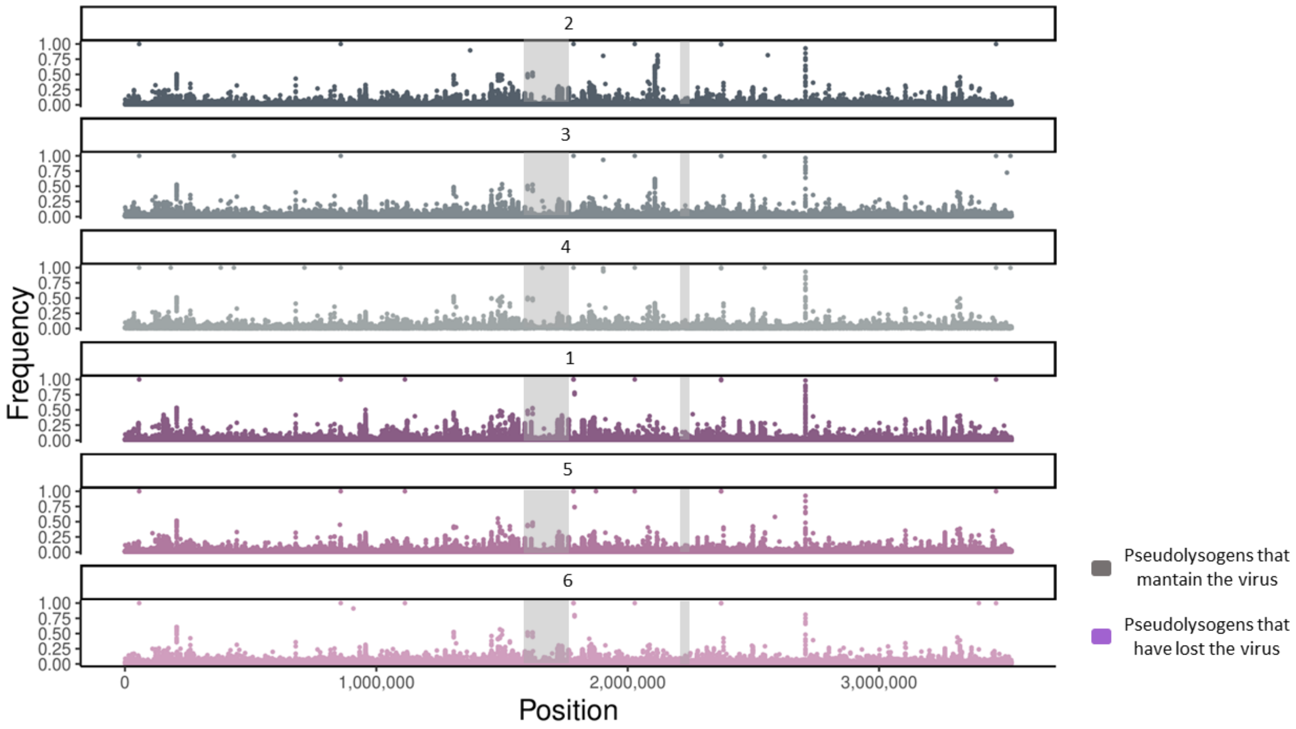


Supplementary Fig. 3. Mutations are not related with virus resistance. Distribution of the mutations (dots) along the chromosome of 3 pseudolysogens that retained the virus the and 3 pseudolysogens that lost the virus. The x-axis represents the length of the chromosome and the y-axis the frequency of each mutation. The first three graphs (top, grey dots) refer to the 3 pseudolysogens that retained the virus (resistants) and the last three (bottom, purple dots) to the 3 pseudolysogens that lost the virus. Hypervariable zones of *Sal. ruber* M1, that correspond to areas of low %GC, are marked in grey

The raw files used for mutation analysis were deposited in the NCBI database with BioProject accession ID PRJNA1136859.

The sequences of the Sanger sequencing of the mutation affecting the sodium/glucose cotransporter can be found below:

>M1+205F

CGCGCTGGGCTCCTTCAGGCTCTTGGCGGCCAGGGCGCGCTGGATGATGTA
CTGGTTGCAGCCCTAGTAGAACAGGTTGGCCACCCACAGGCCCCCAAACAG
CACGCTCAGGCCGGGAAGGAGCTGGTAGGCGTCTTTCAGGGCGCCCTCGTC
GTTGCGGTACATCAGCTAATT

>1+205F

TCCTTCAGGCTCTTGGCGGCCAGGGCGCGCTGGATGATGTACTGGTTGCAGC
CCCAGTAGAACAGGTTGGCCACCCACAGGCCCCCAAACAGCACGCTCAGGC
CGGGAAGGAGCTGGTAGGCGTCTTTCAGGGCGCCCTCGTCGTTGCGGTACA
TCAGCTAG

>2+205F

CTCCTTCAGGCTCTTGGCGGCCAGGGCGCGCTGGATGATGTACTGGTTGCAG
CCCCAGTAGAACAGGTTGGCCACCCACAGGCCCCCAAACAGCACGCTCAGG
CCGGGAAGGAGCTGGTAGGCGTCTTTCAGGGCGCCCTCGTCGTTGCGGTAC
ATCAGCTA

>4+205F

CGCTGGGCTCCTTCAGGCTCTTGGCGGCCAGGGCGCGCTGGATGATGTACTG
GTTGCAGCCCCAGTAGAACAGGTTGGCCACCCACAGGCCCCCAAACAGCAC
GCTCAGGCCGGGAAGGAGCTGGTAGGCGTCTTTCAGGGCGCCCTCGTCGTT
GCGGTACATCAGCTA

>5+205F

GGCGCTGGGCTCCTTCAGGCTCTTGGCGGCCAGGGCGCGCTGGATGATGTA
CTGGTTGCAGCCCCAGTAGAACAGGTTGGCCACCCACAGGCCCCCAAACAG
CACGCTCAGGCCGGGAAGGAGCTGGTAGGCGTCTTTCAGGGCGCCCTCGTC
GTTGCGGTACATCAGCTA

>6+205F

ACCGCCGCTGGGCTCCTTCAGGCTCTTGGCGGCCAGGGCGCGCTGGATGAT
GTACTGGTTGCAGCCCCAGTAGAACAGGTTGGCCACCCACAGGCCCCCAAA
CAGCACGCTCAGGCCGGGAAGGAGCTGGTAGGCGTCTTTCAGGGCGCCCTC
GTCGTTGCGGTACATCAGCTAAT

>7+205F

GGGCCGCTGGGCTCCTTCAGGCTCTTGGCGGCCAGGGCGCGCTGGATGATG
TACTGGTTGCAGCCCCAGTAGAACAGGTTGGCCACCCACAGGCCCCCAAAC
AGCACGCTCAGGCCGGGAAGGAGCTGGTAGGCGTCTTTCAGGGCGCCCTCG
TCGTTGCGGTACATCAGCTAA

>8+205F

CCGCCGCTGGGCTCCTTCAGGCTCTTGGCGGCCAGGGCGCGCTGGATGATGT
ACTGGTTGCAGCCCCAGTAGAACAGGTTGGCCACCCACAGGCCCCCAAACA
GCACGCTCAGGCCGGGAAGGAGCTGGTAGGCGTCTTTCAGGGCGCCCTCGT
CGTTGCGGTACATCAGCTAA

>9+205F

CGCCGCTGGGCTCCTTCAGGCTCTTGGCGGCCAGGGCGCGCTGGATGATGT
ACTGGTTGCAGCCCCAGTAGAACAGGTTGGCCACCCACAGGCCCCCAAACA
GCACGCTCAGGCCGGGAAGGAGCTGGTAGGCGTCTTTCAGGGCGCCCTCGT
CGTTGCGGTACATCAGCTAATTCTAGAAAGAGAGTACTAAAAGAGGAGAGT
GGATGTTCAAGA

>10+205F

CGGCCGCTGGGCTCCTTCAGGCTCTTGGCGGCCAGGGCGCGCTGGATGATG
TACTGGTTGCAGCCCCAGTAGAACAGGTTGGCCACCCACAGGCCCCCAAAC
AGCACGCTCAGGCCGGGAAGGAGCTGGTAGGCGTCTTTCAGGGCGCCCTCG
TCGTTGCGGTACATCAGCTAAT

>11+205F

CCGCCGCTGGGCTCCTTCAGGCTCTTGGCGGCCAGGGCGCGCTGGATGATGT
ACTGGTTGCAGCCCCAGTAGAACAGGTTGGCCACCCACAGGCCCCCAAACA
GCACGCTCAGGCCGGGAAGGAGCTGGTAGGCGTCTTTCAGGGCGCCCTCGT
CGTTGCGGTACATCAGCTAA

>12+205F

AGCCGCCGCTGGGCTCCTTCAGGCTCTTGGCGGCCAGGGCGCGCTGGATGA
TGTACTGGTTGCAGCCCCAGTAGAACAGGTTGGCCACCCACAGGCCCCCAA
ACAGCACGCTCAGGCCGGGAAGGAGCTGGTAGGCGTCTTTCAGGGCGCCCT
CGTCGTTGCGGTACATCAGCTAA

>13+205F

TCCTTCAGGCTCTTGGCGGCCAGGGCGCGCTGGATGATGTACTGGTTGCAGC
CCCAGTAGAACAGGTTGGCCACCCACAGGCCCCCAAACAGCACGCTCAGGC
CGGGAAGGAGCTGGTAGGCGTCTTTCAGGGCGCCCTCGTCGTTGCGGTACA
TCAGCTAATGCGAGAAGAGAGTAGTAAAGTTTGAAGTTGTGCTGCGGT

>14+205F

GGCTCCTTCAGGCTCTTGGCGGCCAGGGCGCGCTGGATGATGTACTGGTTGC
AGCCCCAGTAGAACAGGTTGGCCACCCACAGGCCCCCAAACAGCACGCTCA
GGCCGGGAAGGAGCTGGTAGGCGTCTTTCAGGGCGCCCTCGTCGTTGCGGT
ACATCAGCTAATGATGTTTGTAGAGTGGTGATAGAAGAGGGAGGCGTGTTC

>15+205F

CCGCCGCTGGGCTCCTTCAGGCTCTTGGCGGCCAGGGCGCGCTGGATGATGT
ACTGGTTGCAGCCCCAGTAGAACAGGTTGGCCACCCACAGGCCCCCAAACA
GCACGCTCAGGCCGGGAAGGAGCTGGTAGGCGTCTTTCAGGGCGCCCTCGT
CGTTGCGGTACATCAGCTAATT

>16+205F

AACGGCGCTGGGCTCCTTCAGGCTCTTGGCGGCCAGGGCGCGCTGGATGAT
GTACTGGTTGCAGCCCCAGTAGAACAGGTTGGCCACCCACAGGCCCCCAAA
CAGCACGCTCAGGCCGGGAAGGAGCTGGTAGGCGTCTTTCAGGGCGCCCTC
GTCGTTGCGGTACATCAGCTA

>17+205F

CCGGCGCTGGGCTCCTTCAGGCTCTTGGCGGCCAGGGCGCGCTGGATGATG
TACTGGTTGCAGCCCCAGTAGAACAGGTTGGCCACCCACAGGCCCCCAAAC
AGCACGCTCAGGCCGGGAAGGAGCTGGTAGGCGTCTTTCAGGGCGCCCTCG
TCGTTGCGGTACATCAGCTAA
