## Supplementary material for "Episomal virus maintenance enables bacterial population recovery from infection and virus-bacterial coexistence": Modeling Supplementary Information

#### Model Parameters

| Parameter | Value | Unit |
|---|---|---|
| e, conversion efficiency | 0.75 x $10^{-7}$ | $\mu g$ |
| $\mu_{max}$, maximum growth rate of cells (used to calibrate to experimental curves) | 0.057-0.063 | $h^{-1}$ |
| $R_{in}$, half saturation constant | 8 | $\mu g/ml$ |
| $\lambda_1$, transition rate from $P_e^{[1]}$ to $P_f^{[1]}$ | 1/6.7 | $h^{-1}$ |
| $\eta$, lysis rate | 1/11 | $h^{-1}$ |
| $\beta$, burst size | 10 | |
| $\phi$, adsoprtion rate of free virus into susceptible host | 4 x $10^{-10}$ | $ml/h$ |
| $\sigma$, viral sensitivity to resource levels | 0.06 | |
| $k$, maximum number of superinfected classes | 8 | |
| $\alpha$, selection coefficient for pseudolysogens | 0 | |
| $R_0$, initial resource concentration | 25 | $\mu g/ml$ |

Table 1: Parameters used in ODE model

#### Main model

We develop a nonlinear ODE model to describe the population dynamics of Sal. ruber M1 when under infection by the virus EM1 (Fig. 4a). The host population is denoted as susceptible ($S$), early stage pseudolysogens ($P_e^{[k]}$), and fully formed pseudolysogens ($P_f^{[k]}$), such that the total host population is $N = S + \sum_{i=1}^{i=k}(P_e^{[i]} + P_f^{[i]})$. $P_e^{[k]}$ pseudolysogens are susceptible to superinfection while $P_f^{[k]}$ pseudolysogens are immune to viral adsorption. The variable $k$ indicates the cell MOI, which is the number of viruses infecting a single host cell. Resource consumption by the host population is explicitly modeled through a resource compartment ($R$) that depletes over time. The density of free viruses is denoted by $V$. The model is described by the following system:

$$\dot{R} = -\epsilon\psi(R)(S + \frac{1}{1-\alpha}(\sum_{i=1}^{k} P_e^{[i]} + \sum_{i=1}^{k} P_f^{[i]}))$$

$$\dot{S} = \overbrace{\psi(R)S}^{\text{growth/consumption}} - \overbrace{\phi SV}^{\text{infection}} + \overbrace{2\frac{\psi(R)}{1-\alpha}((\frac{1}{1+1}P_e^{[1]} + \frac{1}{2+1}P_e^{[2]}.....\frac{1}{k+1}P_e^{[k]}) + (\frac{1}{1+1}P_f^{[1]} + \frac{1}{2+1}P_f^{[2]}.....\frac{1}{k+1}P_f^{[k]}))}^{\text{reproductive gain from split inheritance}}$$

$$\dot{P_e^{[1]}} = \overbrace{\phi SV}^{\text{infection}} - \overbrace{\phi P_e^{[1]}V}^{\text{superinfection}} - \overbrace{\lambda_1 P_e^{[1]}}^{\text{transition to full formation}} - \overbrace{\frac{\psi(R)}{1-\alpha}P_e^{[1]}}^{\text{reproductive loss}} + \overbrace{2\frac{\psi(R)}{1-\alpha}(\frac{1}{1+1}P_e^{[1]} + \frac{1}{2+1}P_e^{[2]} + .....\frac{1}{k+1}P_e^{[k]})}^{\text{reproductive gain}}$$

$$\dot{P_e^{[2]}} = \overbrace{\phi P_e^{[1]}V}^{\text{superinfection in}} - \overbrace{\phi P_e^{[2]}V}^{\text{superinfection out}} - \lambda_2 P_e^{[2]} - \frac{\psi(R)}{1-\alpha}P_e^{[2]} + 2\frac{\psi(R)}{1-\alpha}(\frac{1}{2+1}P_e^{[2]} + \frac{1}{3+1}P_e^{[3]} + .....\frac{1}{k+1}P_e^{[k]})$$

$$\vdots$$

$$\dot{P_e^{[k']}} = \phi P_e^{[k'-1]}V - \phi P_e^{[k']}V - \lambda_k' P_e^{[k']} - \frac{\psi(R)}{1-\alpha}P_e^{[k']} + 2\frac{\psi(R)}{1-\alpha}(\sum_{i=k'}^{k}\frac{1}{i+1}P_e^{[i]})$$

$$\vdots$$

$$\dot{P_e^{[k]}} = \phi P_e^{[k-1]} V - \lambda_k P_e^{[k]} - \frac{\psi(R)}{1-\alpha} P_e^{[k]} + 2\frac{\psi(R)}{1-\alpha}(\frac{1}{k+1} P_e^{[k]})$$

$$\dot{P_f^{[1]}} = \overbrace{\lambda_1 P_e^{[1]}}^{\text{transition to full formation}} - \overbrace{\frac{\psi(R)}{1-\alpha} P_f^{[1]}}^{\text{reproductive loss}} + \overbrace{2\frac{\psi(R)}{1-\alpha}(\frac{1}{1+1} P_f^{[1]} + \frac{1}{2+1} P_f^{[2]} + \ldots . . \frac{1}{k+1} P_f^{[k]})}^{\text{reproductive gain}} - \overbrace{\eta\gamma P_f^{[1]}}^{\text{lysis}}$$

$$\dot{P_f^{[2]}} = \lambda_2 P_e^{[2]} - \frac{\psi(R)}{1-\alpha} P_f^{[2]} + 2\frac{\psi(R)}{1-\alpha}(\frac{1}{2+1} P_f^{[2]} + \frac{1}{3+1} P_f^{[3]} + \ldots . . \frac{1}{k+1} P_f^{[k]}) - \eta\gamma P_f^{[2]}$$

$$\vdots$$

$$\dot{P_f^{[k]}} = \lambda_k P_e^{[k]} - \frac{\psi(R)}{1-\alpha} P_f^{[k]} + 2\frac{\psi(R)}{1-\alpha}(\frac{1}{k+1} P_f^{[k]}) - \eta\gamma P_f^{[k]}$$

$$\dot{V} = \overbrace{\beta\gamma\eta \sum_{i=1}^{k} P_f^{[i]}}^{\text{burst}} - \overbrace{\phi(S + \sum_{i=1}^{k-1} P_e^{[i]})V}^{\text{total infection}}$$

Given the resource-explicit dynamics, $\psi(R) = \mu_{max}(R/(R + R_{in}))$ is the Monod equation, where $\mu_{max}$ is the maximal growth rate and $R_{in}$ is the resource concentration at which the growth rate is half its maximum. Parameter $e$ is the conversion efficiency, $\alpha$ is the selective advantage in reproductive output that pseudolysogens (E) have over susceptible cells (S).

Function $\gamma$ modulates lysis depending on the resource level and is equal to $(\frac{R}{R+\sigma R_{in}})/(\frac{R_0}{R_0+\sigma R_{in}})$, where $R_0$ is the initial resource concentration at t=0 and $\sigma$ is the viral sensitivity to starvation. Additionally, parameter $\phi$ is the adsorption rate of free viruses into host cells, $\lambda_k$ is the transition rate from an early stage pseudolysogen with $k$ viral genomes to a fully formed pseudolysogen with $k$ viral genomes, $\eta$ is the lysis rate, and $\beta$ is the burst size. We use a multistage infection model where the latent period is equal to $1/\lambda_k + 1/\eta$. We make the transition rate $\lambda_k$ decrease as a function of $k$, which results in the latent period increasing as a function of $k$ (Fig. S1). Mechanistically, this has the important feature of delaying lysis upon the adsorption of additional viruses.

### Key Model Features

As we identified EM1 to not have a lysogenic infection pathway, we initially used a purely lytic ODE model (Supplement: Pure lytic) to simulate the expected population dynamics of Sal. ruber M1 under infection at different bulk MOIs. The simulated results (Figure S4) highlight that in a purely lytic system, the host population should eventually crash regardless of MOI and that the higher the MOI, the earlier and sharper the expected decline in the host population should be. Evidently, the simulated results from the pure lytic model are vastly different to the experiment results observed (Fig 3c). We break down the key experimental dynamics that the pure lytic model fails to capture, with a focus on three key features:

1. The M1 populations infected at MOI 0.1 and 0.01 do not experience an observable population dip, highlighting a lack of expected killing
2. The M1 host population recovers to a higher population when infected at MOI 10 when compared to MOI 5 and MOI 1
3. When infected at MOI 10, the M1 population does not crash immediately upon infection and maintains significant transient growth post infection.

##### Host lysis is delayed when available resources are depleted

Studies have shown that in some virus-host systems (e.g, *E.coli* - T4), a lack of expected killing is observed when resources get depleted enough to induce starvation in the infected host (40,41). As feature 1 highlights such a lack of expected killing in M1-EM1, we iterated upon the obligately lytic model but this time making lysis events sensitive to the level of resources present in the culture. Full details of the model specifications are found in Supplement: *Pure resource dependent lysis inhibition*. To summarize, we introduce a function $\gamma = (\frac{R}{R+\sigma R_{in}})/(\frac{R_0}{R_0+\sigma R_{in}})$, where $\sigma$ = viral sensitivity to host starvation. By creating an effective lysis rate $\eta\gamma$, we allow for lysis rates to be independent when resources are abundant ($R \gg \sigma R_{in}$) but reduce as a function of resources as resources are depleted through consumption.

The resulting simulated population dynamics of Sal. ruber M1 (Figure S5a) shows the populations infected at MOI 0.1 and 0.01 not crashing as they did in the purely lytic simulations (Figure S4), capturing feature 1 in the process. However, the simulated dynamics seen at the higher bulk MOIs still show the populations crashing in similar fashion to the simulated pure lytic dynamics, thereby failing to capture features 2 and 3. This failure can be explained by tracking the simulated resource levels over time(Fig. S5b), where we find that the fixed resource well is minimally depleted in the cultures infected at higher MOIs when compared to the populations infected at the lower MOIs. This implies that there is significant killing of the simulated host populations at higher bulk MOIs and therefore not enough host resource consumption to trigger resource dependent lysis inhibition.

**Infected host lysis is delayed upon adsorption of additional viruses**

Studies of lysis inhibition have shown that the adsorption of additional bacteriophage into a host can extend the infection's latent period, inhibiting/delaying lysis in the process (42,43). As we have already shown that multiple viruses can infect a single host cell in our system, we decided to explore whether the addition of superinfection induced lysis inhibition can allow our simulation to capture more experimental features. We iterate on the previous model that solely implemented resource dependent lysis inhibition by now allowing for the individual simulated Sal. ruber M1 host cells to be superinfected by multiple viruses (Figure S6). Full details of the model specifications are found in Supplement: *Superinfection induced + resource dependent lysis inhibition*. It is important to note that in this iteration of the model we do not allow for limitless superinfection to occur. By varying the number of viruses that can infect a single host in our model, we found that limiting the maximum number of viruses (k) that can infect a single host to eight qualitatively produced simulated adsorption curves that matched the experimental curves observed at MOI 10 (Extended Data Fig. 3). Therefore for all the following results, we set k = 8.

The main feature of this iteration of the model is that the latent period, which is the elapsed time from initial viral adsorption to host lysis, increases as a function of superinfection (Fig S1). Mechanistically this means that the more viruses infecting a single host, the longer the host survives before eventual lysis. The addition of superinfection induced lysis inhibition alongside resource dependent lysis inhibition prevents the populations infected at bulk MOI 1, 5 and 10 from crashing as seen in the previous iteration of the model. Furthermore, the addition captures the dynamics seen in feature 2, where the simulated population infected at bulk MOI 10 has better final outcomes than the populations infected at bulk MOI 5 and MOI 1.

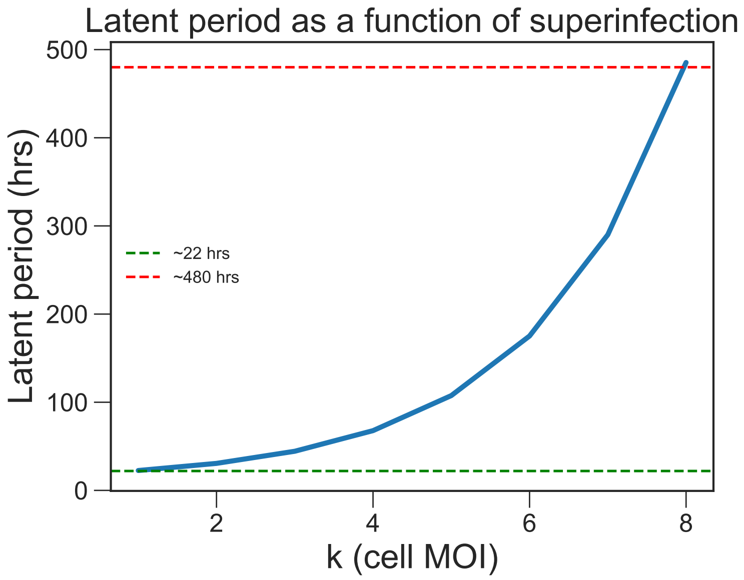


Figure S1: Graph of how the model varies the latent period as a function of superinfection: latent period = $1/\lambda_k$ $+1/\eta$ , where $\lambda_k = \lambda_1/(a)^{k-1}$. Threshold maximum latent period $\sim$ 480 hrs with $a = 1.7$, $\lambda_1 = 1/6.7$ (subscript denotes cell MOI $k$) and $\eta = 1/11$. The latent period without superinfection (22 hours) was experimentally measured using a one-step growth curve (Methods: One step growth analysis). Note that a linear increase in latent period as a function of superinfection was also attempted found to be unsatisfactory, thereby justifying the use of a non linear increasing function.

**Cell division of superinfected pseudolysogens allows for transient host population growth and vertical virus transmission**

While successful in capturing feature 1 and 2, lysis inhibition does not explain how the same population infected at MOI 10 undergoes significant transient growth post viral inoculation as noted in feature 3. This transient growth implies that there is persistent cell division of the infected pseudolysogens even under significant net infection. Most studies that explore pseudolysogen cell division suggest that the viral genome is asymmetrically passed down to only one of the daughter cells; however, these studies only consider single infections of pseudolysogens (23, 24) while our model allows for multiple viral genomes to exist within a single host cell through superinfection. Therefore to model the inheritance of multiple viral genomes upon pseudolysogen cell division, we iterate upon our lysis inhibition model and propose a theoretical model of superinfection-dependent asymmetric cell division. We propose two variants of the model in the 'split inheritance' and the 'prioritized inheritance' frameworks (Supplement *Viral passage in superinfected pseudolysogens*). We will focus on the results given from the 'split inheritance' framework, where the superinfected viral load of the parent is randomly split amongst its daughter cells, allowing for both daughter cells to get vertically infected. Pl. Full details of the model specification are found in Supplement *Main Model*.

It is important to note that we separate the pseudolysogens into early stage pseudolysogens ($P_e^k$) and fully formed pseudolysogens ($P_f^k$). As we know that superinfection is possible insofar as it precedes the development of surface level resistance, we separate the pseudolysogens to allow early stage pseudolysogens ($P_e^k$) to get superinfected while making fully formed pseudolysogens ($P_f^k$) resistant to further viral infection at the surface level. Implementing pseudolysogen cell division through the 'split inheritance' framework proved to be the final piece in our iterating model puzzle. The experimental dynamics, defined by the aforementioned three key features, are successfully captured by the simulated results (Fig. 5a). Deeper analysis into the simulated results reveals that while there is more net infection occurring at MOI 10 (Fig. S2a), the infected pseudolysogens are skewed towards being more superinfected in their early stages (Fig. S2b). These superinfected pseudolysogens in turn survive longer than their less superinfected counterparts due to their longer latent periods (Fig. S1). This leads to better final retention of the host population when infected at MOI 10 than at MOI 5 or 1, and the unintuitive outcome that more infection can lead to larger surviving final host populations. We hypothesize that this superinfected driven retention seen in our simulations is reflected in reality as well, helping to explain why superinfection induced lysis inhibition captures feature 2. Furthermore, we hypothesize that the presence highly superinfected pseudolysogens with long lifespans that continue to cell divide even upon infection can be directly correlated to the transient growth observed post viral inoculation (66 hours) at MOI 10; thereby capturing feature 3 and wrapping up the final successful iteration of the model seen in Fig. 5a.

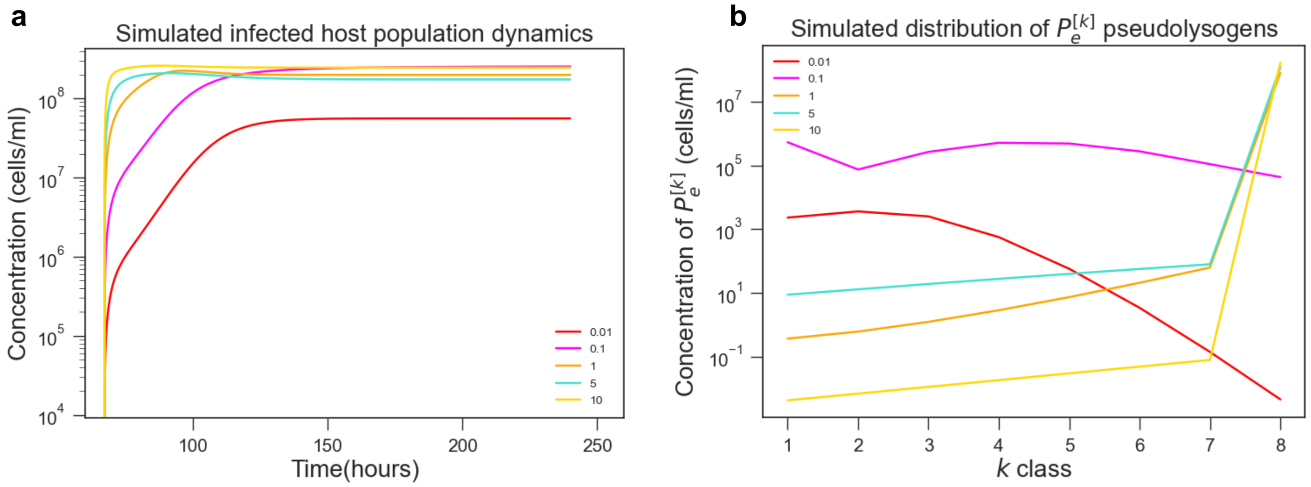


Figure S2: **a**, Simulated population dynamics of the total infected $\sum_{i=1}^{i=k}(P_e^{[i]} + P_f^{[i]})$ hosts. **b** Distribution of early stage pseudolysogens ($P_e^k$) hosts across the superinfection classes (k). The higher the k class of the $P_e^k$ pseudolysogen, the more the pseudolysogen's eventual lysis is delayed due to its longer latent period

### Viral passage in superinfected pseudolysogens

For this study we hypothesize that the supeinfected viral load of the parent pseudolysogen is randomly distributed to the daughter cells upon cell division in a process we name split inheritance. However, as stated earlier past work on pseudolysogeny has stated that upon cell division, one of the daughter cells is virus-free. Therefore we explore two different theoretical models of viral passage in pseudolysogens: split inheritance and prioritized inheritance, where prioritized inheritance depicts the entirety of the viral genomes present in the parent being passed on to a single daughter cell, leaving one of the daughter cells virus-free. Implementing both models produce the same population dynamics for Sal. Ruber M1 (Fig. S3) as both model pseudolysogen cell division, they key model feature that is needed to explain transient population growth at high MOIs. However, the implications of both models on the viral presence in the final population is drastically different.

In the split inheritance framework, there are $k+1$ (where $k$ is the cell MOI of the parent) configurations of daughter cell pairs that can be produced, and $2k+2$ possible daughter cells. Out of the $2k+2$ daughter cells, $2k$ will be pseudoslysogens as only the daughter cell pairs (k,0) and (0,k) will produce virus free daughters. This implies that $\frac{k}{k+1}$ of the next generation of children of pseudolysogens will themselves be pseudolysogens. Furthermore, a $P^k$ pseudolysogen (where the following dynamics of $P^k$ reflect the dynamics of both $P_e^k$ and $P_f^k$) has a $2/(k+1)$ probability of producing a $P^{k'}$ pseudolysogen where $k$' $\leq k$. This is implemented in the ODE model through the denoted 'split inheritance' terms, where there is 'reproductive gain' from the $2/(k+1)$ daughters of all the $P^k$ classes and reproductive loss from all the daughter cells produced by $P^{k'}$. The repopulation of $P^k$ pseudolysogens, alongside the abundance and longevity of the highly superinfected pseudolysogens produced when total infection is high at MOI 10 (Fig. S2b), can be attributed to the significant transient growth post infection observed through feature 3.

This repopulation of the $P^k$ classes in the split inheritance framework can also be attributed to the maintained viral presence, as the viral load is vertically transmitted through cell division. With prioritized inheritance however, the viral load will only passage along one branch of the cell division binary tree. This implies that after $t$ generations, only a single daughter cell out of the entire population ($2^t$) will contain the entirety of the original parent's viral load $k$. This leads to no repopulation of the $P^k$ classes, and only repopulation of the virus-free $S$ class through pseudolysogen cell division (Section: prioritized inheritance Model). Therefore, the proportion of the host population that is infected dilutes significantly quicker in the prioritized inheritance framework than in the split inheritance framework. Single cell analysis is required to confirm which framework is present in the M1-EM1 system, which is beyond the scope of this study. However akin to the random segregation of multiple intracellular plasmids, it seems unlikely that there would be a bias towards one daughter cell getting the entirety of the viral genome load from the parent, and therefore the main ODE model for this study uses split inheritance.

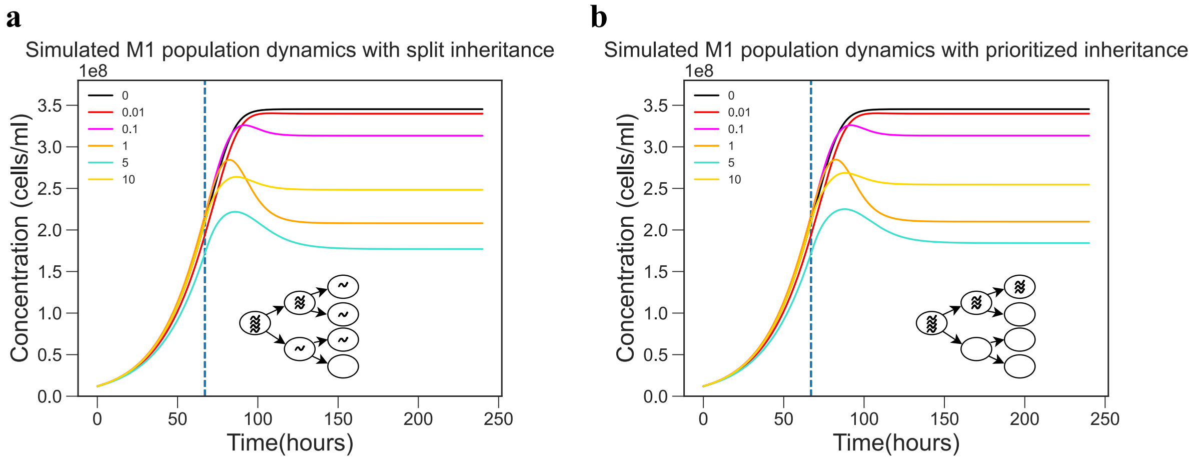


Figure S3: **a**, Simulated population dynamics when implementing split inheritance, with an inset schematic showing an example of split inheritance cell division. **b**, Simulated population dynamics when implementing prioritized inheritance, with an inset schematic showing an exmple of prioritized inheritance cell division. (refer to *Supplement: Prioritized inheritance model*)

### Prioritized inheritance model

$$\dot{R} = -\epsilon\psi(R)(S + \frac{1}{1-\alpha}(\sum_{i=1}^{k} P_e^{[i]} + \sum_{i=1}^{k} P_f^{[i]}))$$

$$\dot{S} = \overbrace{\psi(R)S}^{\text{growth/consumption}} - \overbrace{\phi SV}^{\text{infection}} + \overbrace{\frac{\psi(R)}{1-\alpha}(\sum_{i=1}^{k} P_e^{[i]} + \sum_{i=1}^{k} P_f^{[i]})}^{\text{reproductive gain from prioritized inheritance}}$$

$$\dot{P_e^{[1]}} = \overbrace{\phi SV}^{\text{infection}} - \overbrace{\phi P_e^{[1]}V}^{\text{superinfection}} - \overbrace{\lambda_1 P_e^{[1]}}^{\text{transition to full formation}}$$

$$\dot{P_e^{[2]}} = \overbrace{\phi P_e^{[1]}V}^{\text{superinfection in}} - \overbrace{\phi P_e^{[2]}V}^{\text{superinfection out}} - \lambda_2 P_e^{[2]}$$

$$\vdots$$

$$\dot{P_e^{[k']}} = \phi P_e^{[k'-1]}V - \phi P_e^{[k']}V - \lambda_k' P_e^{[k']}$$

$$\vdots$$

$$\dot{P_e^{[k]}} = \phi P_e^{[k-1]}V - \lambda_k P_e^{[k]}$$

$$\dot{P_f^{[1]}} = \overbrace{\lambda_1 P_e^{[1]}}^{\text{transition to full formation}} - \overbrace{\eta\gamma P_f^{[1]}}^{\text{lysis}}$$

$$\dot{P_f^{[2]}} = \lambda_2 P_e^{[2]} - \eta\gamma P_f^{[2]}$$

$$\vdots$$

$$\dot{P_f^{[k]}} = \lambda_k P_e^{[k]} - \eta\gamma P_f^{[k]}$$

$$\dot{V} = \overbrace{\beta\gamma\eta \sum_{i=1}^{k} P_f^{[i]}}^{\text{burst}} - \overbrace{\phi(S + \sum_{i=1}^{k} P_e^{[i]})V}^{\text{total infection}}$$

### Subsets of main model with corresponding dynamics

#### Pure lytic

$$\dot{R} = \overbrace{-\epsilon\psi(R)S}^{\text{consumption}}$$

$$\dot{S} = \overbrace{\psi(R)S}^{\text{growth}} - \overbrace{\phi SV}^{\text{infection}}$$

$$\dot{E} = \phi SV - \overbrace{\lambda E}^{\text{commitment transition to full formation}}$$

$$\dot{I} = \lambda E - \overbrace{\eta I}^{\text{lysis}}$$

$$\dot{V} = \overbrace{\beta\eta}^{\text{burst}} - \overbrace{\phi(S + E + I)V}^{\text{total infection}}$$

where $\psi(R) = \frac{\mu_{max} R}{R + R_{in}}$

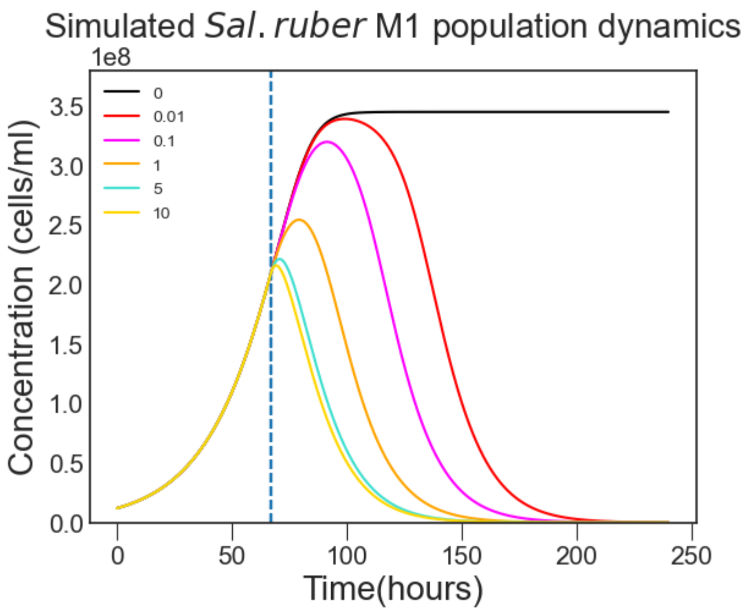


Figure S4: Simulated population dynamics with just pure resource dependent lysis inhibition

#### Pure resource dependent lysis inhibition

$$\dot{R} = -\epsilon\psi(R)S$$

$$\dot{S} = \overbrace{\psi(R)S}^{\text{growth/consumption}} - \overbrace{\phi SV}^{\text{infection}}$$

$$\dot{P}_e = \overbrace{\phi SV}^{\text{infection}} - \overbrace{\phi P_e^{[1]} V}^{\text{superinfection}} - \overbrace{\lambda P_e^{[1]}}^{\text{transition to full formation}}$$

$$\dot{P}_f = \overbrace{\lambda P_e^{[1]}}^{\text{transition to full formation}} \overbrace{- \eta\gamma P_f^{[1]}}^{\text{lysis}}$$

$$\dot{V} = \overbrace{\beta\gamma\eta P_f}^{\text{burst}} - \overbrace{\phi(S + P_e)V}^{\text{total infection}}$$

where $\gamma = (\frac{R}{R+\sigma R_{in}})/(\frac{R_0}{R_0+\sigma R_{in}})$ and $\sigma =$ viral sensitivity to host starvation

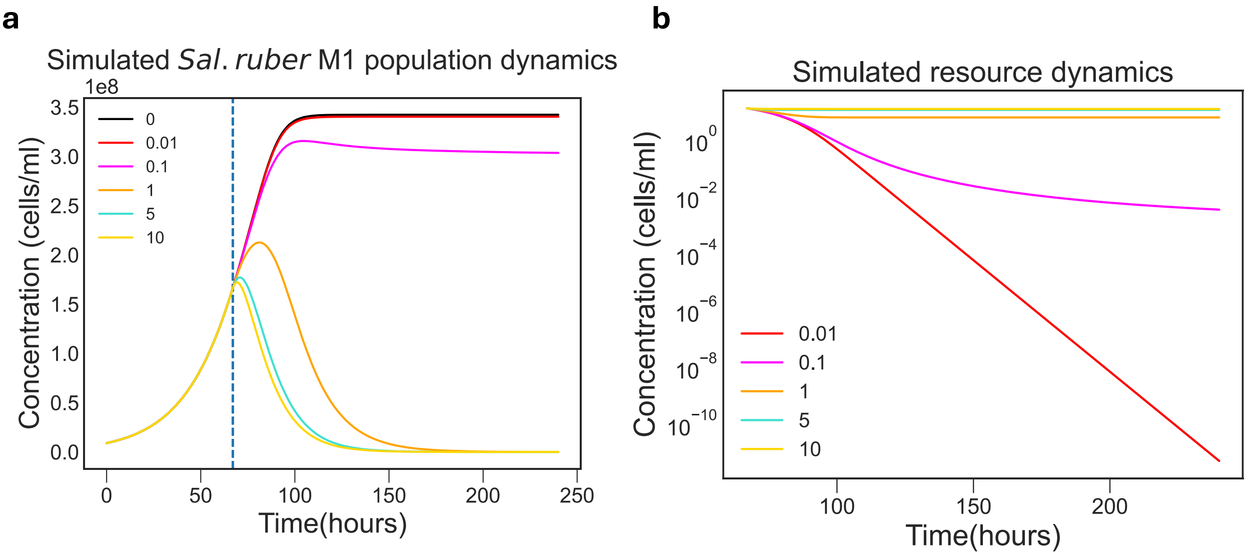


‘

Figure S5: Expected simulated results when solely implementing resource dependent lysis inhibition. **a**, Simulated Sal. ruber M1 population dynamics upon infection by virus EM1 with resource dependent lysis inhibition (Supplement: Pure resource dependent lysis inhibition). **b**, Simulated resource depletion over time. It is important to note that resource depletion is significantly less at high MOIs (5,10), making pure resource dependent lysis inhibition ineffective with high levels of infection

#### Superinfection induced + resource dependent lysis inhibition

$$\dot{R} = -\epsilon\psi(R)(S + \frac{1}{1-\alpha}(\sum_{i=1}^{k} P_e^{[i]} + \sum_{i=1}^{k} P_f^{[i]}))$$

$$\dot{S} = \overbrace{\psi(R)S}^{\text{growth/consumption}} - \overbrace{\phi SV}^{\text{infection}}$$

$$\dot{P_e^{[1]}} = \overbrace{\phi SV}^{\text{infection}} - \overbrace{\phi P_e^{[1]} V}^{\text{superinfection}} - \overbrace{\lambda_1 P_e^{[1]}}^{\text{transition to full formation}}$$

$$\dot{P_e^{[2]}} = \overbrace{\phi P_e^{[1]} V}^{\text{superinfection in}} - \overbrace{\phi P_e^{[2]} V}^{\text{superinfection out}} - \lambda_2 P_e^{[2]}$$

$$\vdots$$

$$\dot{P_e^{[k']}} = \phi P_e^{[k'-1]} V - \phi P_e^{[k']} V - \lambda'_k P_e^{[k']}$$

$$\vdots$$

$$\dot{P_e^{[k]}} = \phi P_e^{[k-1]} V - \lambda_k P_e^{[k]}$$

$$\dot{P_f^{[1]}} = \overbrace{\lambda_1 P_e^{[1]}}^{\text{transition to full formation}} - \overbrace{\eta\gamma P_f^{[1]}}^{\text{lysis}}$$

$$\dot{P_f^{[2]}} = \lambda_2 P_e^{[2]} - \eta\gamma P_f^{[2]}$$

$$\vdots$$

$$\dot{P_f^{[k]}} = \lambda_k P_e^{[k]} - \eta\gamma P_f^{[k]}$$

$$\dot{V} = \overbrace{\beta\gamma\eta\sum_{i=1}^{k} P_f^{[i]}}^{\text{burst}} - \overbrace{\phi(S + \sum_{i=1}^{k} P_e^{[i]})V}^{\text{total infection}}$$

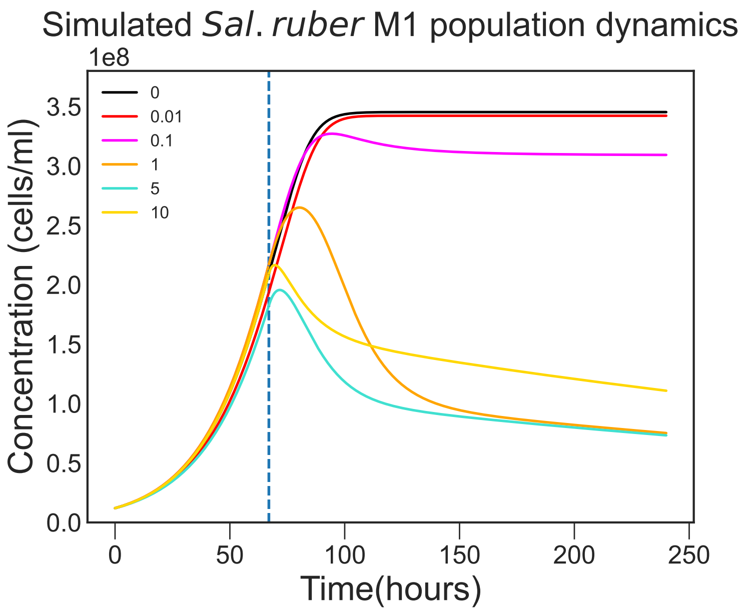


Figure S6: Simulated population dynamics with just superinfected induced lysis inhibition and resource dependent lysis inhibition

#### Pure superinfection induced lysis inhibition

$$\dot{R} = -\epsilon\psi(R)(S + \frac{1}{1-\alpha}(\sum_{i=1}^{k} P_e^{[i]} + \sum_{i=1}^{k} P_f^{[i]}))$$

$$\dot{S} = \overbrace{\psi(R)S}^{\text{growth/consumption}} - \overbrace{\phi SV}^{\text{infection}}$$

$$\dot{P_e^{[1]}} = \overbrace{\phi SV}^{\text{infection}} - \overbrace{\phi P_e^{[1]} V}^{\text{superinfection}} - \overbrace{\lambda_1 P_e^{[1]}}^{\text{transition to full formation}}$$

$$\dot{P_e^{[2]}} = \overbrace{\phi P_e^{[1]} V}^{\text{superinfection in}} - \overbrace{\phi P_e^{[2]} V}^{\text{superinfection out}} - \lambda_2 P_e^{[2]}$$

$$\vdots$$

$$\dot{P_e^{[k']}} = \phi P_e^{[k'-1]} V - \phi P_e^{[k']} V - \lambda_k' P_e^{[k']}$$

$$\vdots$$

$$\dot{P_e^{[k]}} = \phi P_e^{[k-1]} V - \lambda_k P_e^{[k]}$$

$$\dot{P_f^{[1]}} = \overbrace{\lambda_1 P_e^{[1]}}^{\text{transition to full formation}} - \overbrace{\eta P_f^{[1]}}^{\text{lysis}}$$

$$\dot{P_f^{[2]}} = \lambda_2 P_e^{[2]} - \eta P_f^{[2]}$$

$$\vdots$$

$$\dot{P_f^{[k]}} = \lambda_k P_e^{[k]} - \eta P_f^{[k]}$$

$$\dot{V} = \overbrace{\beta\eta\sum_{i=1}^{k} P_f^{[i]}}^{\text{burst}} - \overbrace{\phi(S + \sum_{i=1}^{k} P_e^{[i]})V}^{\text{total infection}}$$

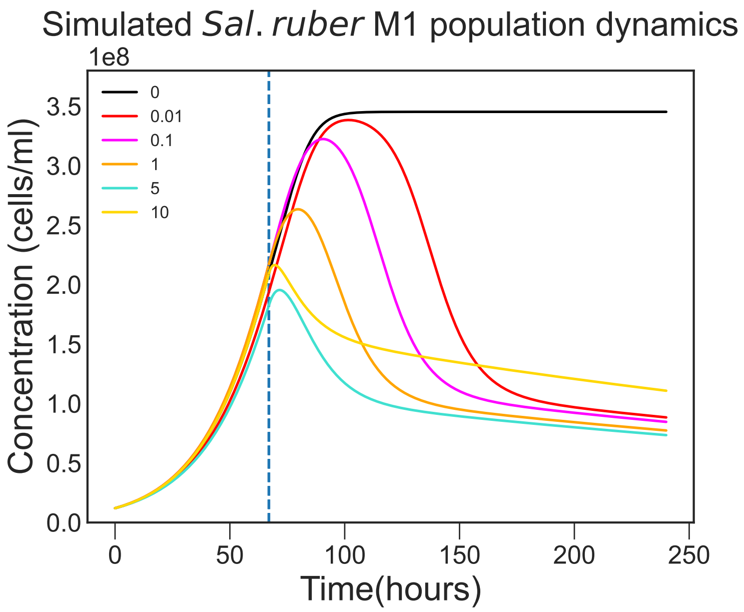


Figure S7: Simulated population dynamics with just pure superinfection induced lysis inhibition

#### Pure pseudolysogenic cell division (split inheritance)

$$\dot{R} = -\epsilon\psi(R)S$$

$$\dot{S} = \overbrace{\psi(R)S}^{\text{growth/consumption}} - \overbrace{\phi SV}^{\text{infection}} + \overbrace{\frac{\psi(R)}{1-\alpha}(P_e + P_f)}^{\text{reproductive gain from split inheritance}}$$

$$\dot{P}_e = \overbrace{\phi SV}^{\text{infection}} - \overbrace{\phi P_e^{[1]}V}^{\text{superinfection}} - \overbrace{\lambda P_e^{[1]}}^{\text{transition to full formation}}$$

$$\dot{P}_f = \overbrace{\lambda P_e^{[1]}}^{\text{transition to full formation}} \overbrace{-\eta P_f^{[1]}}^{\text{lysis}}$$

$$\dot{V} = \overbrace{\beta\eta P_f}^{\text{burst}} - \overbrace{\phi(S + P_e)V}^{\text{total infection}}$$

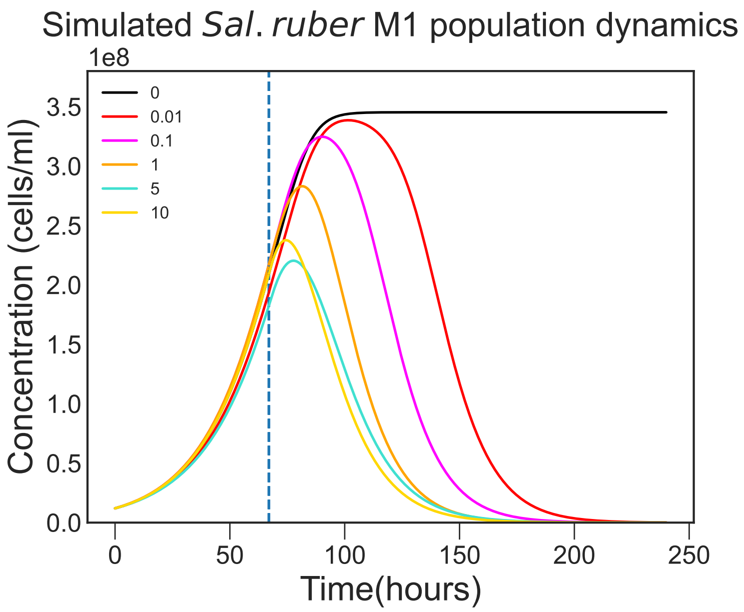


Figure S8: Simulated population dynamics with just pure pseudolysogeny

#### Superinfection induced lysis inhibition + pseudolysogenic cell division (split inheritance)

$$\dot{R} = -\epsilon\psi(R)(S + \frac{1}{1-\alpha}(\sum_{i=1}^{k} P_e^{[i]} + \sum_{i=1}^{k} P_f^{[i]}))$$

$$\dot{S} = \overbrace{\psi(R)S}^{\text{growth/consumption}} - \overbrace{\phi SV}^{\text{infection}} + \overbrace{2\frac{\psi(R)}{1-\alpha}((\frac{1}{1+1}P_e^{[1]} + \frac{1}{2+1}P_e^{[2]}.....\frac{1}{k+1}P_e^{[k]}) + (\frac{1}{1+1}P_f^{[1]} + \frac{1}{2+1}P_f^{[2]}.....\frac{1}{k+1}P_f^{[k]}))}^{\text{reproductive gain from split inheritance}}$$

$$\dot{P_e^{[1]}} = \overbrace{\phi SV}^{\text{infection}} - \overbrace{\phi P_e^{[1]} V}^{\text{superinfection}} - \overbrace{\lambda_1 P_e^{[1]}}^{\text{transition to full formation}} - \overbrace{\frac{\psi(R)}{1-\alpha}P_e^{[1]}}^{\text{reproductive loss}} + \overbrace{2\frac{\psi(R)}{1-\alpha}(\frac{1}{1+1}P_e^{[1]} + \frac{1}{2+1}P_e^{[2]} + .....\frac{1}{k+1}P_e^{[k]})}^{\text{reproductive gain}}$$

$$\dot{P_e^{[2]}} = \overbrace{\phi P_e^{[1]} V}^{\text{superinfection in}} - \overbrace{\phi P_e^{[2]} V}^{\text{superinfection out}} - \lambda_2 P_e^{[2]} - \frac{\psi(R)}{1-\alpha}P_e^{[2]} + 2\frac{\psi(R)}{1-\alpha}(\frac{1}{2+1}P_e^{[2]} + \frac{1}{3+1}P_e^{[3]} + .....\frac{1}{k+1}P_e^{[k]})$$

$$\vdots$$

$$\dot{P_e^{[k']}} = \phi P_e^{[k'-1]} V - \phi P_e^{[k']} V - \lambda'_k P_e^{[k']} - \frac{\psi(R)}{1-\alpha}P_e^{[k']} + 2\frac{\psi(R)}{1-\alpha}(\sum_{i=k'}^{k}\frac{1}{i+1}P_e^{[i]})$$

$$\vdots$$

$$\dot{P_e^{[k]}} = \phi P_e^{[k-1]} V - \lambda_k P_e^{[k]} - \frac{\psi(R)}{1-\alpha}P_e^{[k]} + 2\frac{\psi(R)}{1-\alpha}(\frac{1}{k+1}P_e^{[k]})$$

$$\dot{P_f^{[1]}} = \overbrace{\lambda_1 P_e^{[1]}}^{\text{transition to full formation}} - \overbrace{\frac{\psi(R)}{1-\alpha}P_f^{[1]}}^{\text{reproductive loss}} + \overbrace{2\frac{\psi(R)}{1-\alpha}(\frac{1}{1+1}P_f^{[1]} + \frac{1}{2+1}P_f^{[2]} + .....\frac{1}{k+1}P_f^{[k]})}^{\text{reproductive gain}} - \overbrace{\eta P_f^{[1]}}^{\text{lysis}}$$

$$\dot{P_f^{[2]}} = \lambda_2 P_e^{[2]} - \frac{\psi(R)}{1-\alpha}P_f^{[2]} + 2\frac{\psi(R)}{1-\alpha}(\frac{1}{2+1}P_f^{[2]} + \frac{1}{3+1}P_f^{[3]} + .....\frac{1}{k+1}P_f^{[k]}) - \eta P_f^{[2]}$$

$$\vdots$$

$$\dot{P_f^{[k]}} = \lambda_k P_e^{[k]} - \frac{\psi(R)}{1-\alpha}P_f^{[k]} + 2\frac{\psi(R)}{1-\alpha}(\frac{1}{k+1}P_f^{[k]}) - \eta P_f^{[k]}$$

$$\dot{V} = \overbrace{\beta\eta\sum_{i=1}^{k} P_f^{[i]}}^{\text{burst}} - \overbrace{\phi(S + \sum_{i=1}^{k-1} P_e^{[i]})V}^{\text{total infection}}$$

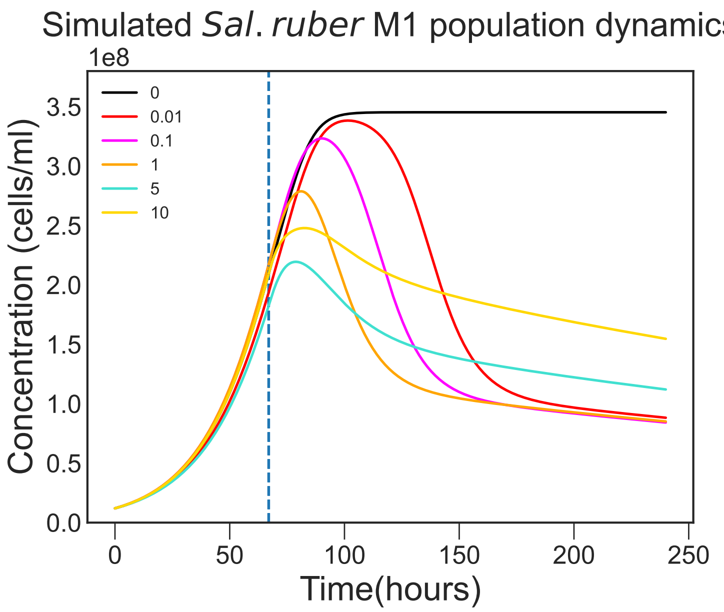


Figure S9: Simulated population dynamics with just superinfection induced lysis inhibition and pseudolysogeny (split inheritance)

##### Resource limited lysis inhibition + pseudolysogenic cell division (split inheritance)

$$\dot{R} = -\epsilon\psi(R)S$$

$$\dot{S} = \overbrace{\psi(R)S}^{\text{growth/consumption}} - \overbrace{\phi SV}^{\text{infection}} + \overbrace{\frac{\psi(R)}{1-\alpha}(P_e + P_f)}^{\text{reproductive gain from split inheritance}}$$

$$\dot{P}_e = \overbrace{\phi SV}^{\text{infection}} - \overbrace{\phi P_e^{[1]} V}^{\text{superinfection}} - \overbrace{\lambda P_e^{[1]}}^{\text{transition to full formation}}$$

$$\dot{P}_f = \overbrace{\lambda P_e^{[1]}}^{\text{transition to full formation}} - \overbrace{\eta\gamma P_f^{[1]}}^{\text{lysis}}$$

$$\dot{V} = \overbrace{\beta\eta\gamma P_f}^{\text{burst}} - \overbrace{\phi(S + P_e)V}^{\text{total infection}}$$

#### Adsoprtion rate calculation for M1-EM1 system

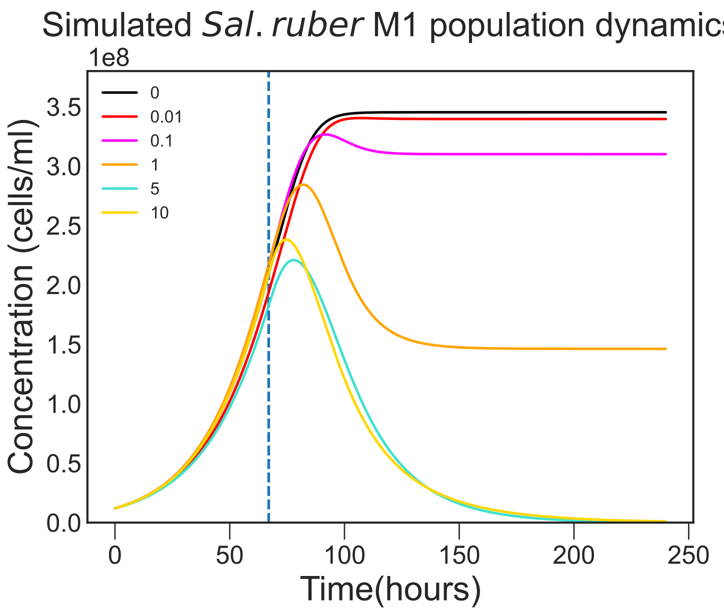


Figure S10: Simulated population dynamics with just resource limited lysis inhibition and pseudolysogeny (split inheritance)

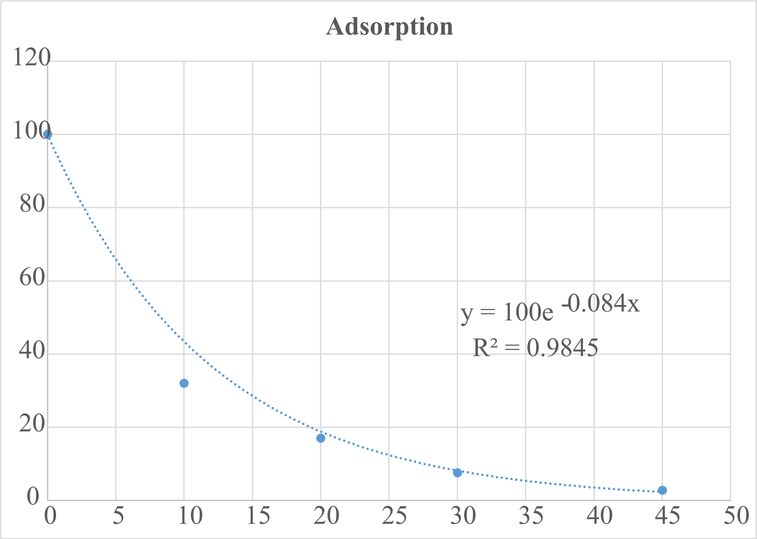


Figure S11: Solving $\dot{V} = -\phi SV$ gives $V = V_0 e^{-\phi St}$. The one step experiment was done with an initial $S = 2$ x $10^8 cells/ml$ and we can assume that in 60 minutes there is negligible killing (latent period = 21-22 hours). Therefore, we equate (2 x $10^8$)$\phi$ with the fitted exponent 0.084 to get $\phi \approx 4$ x $10^{-10} ml/h$

#### Revival

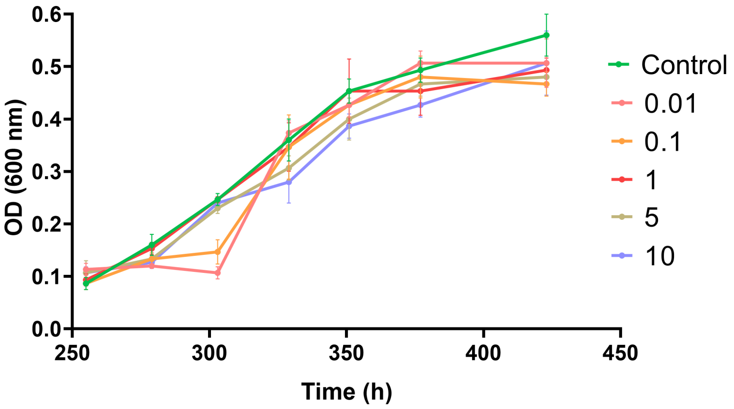


Figure S12: Population dynamics upon resource reintroduction after MOI experiment from Figure 3a. The early stationary phase observed at MOI 0.01 and 0.1 (250-300 min) can be attributed to lysis events. These lysis events correspond to killing off the infected host cells that were starved and therefore lysis inhibited due to resource limitations.

#### Sal.ruber M1 wild-type and 1R (pseudolysogen) growth comparison

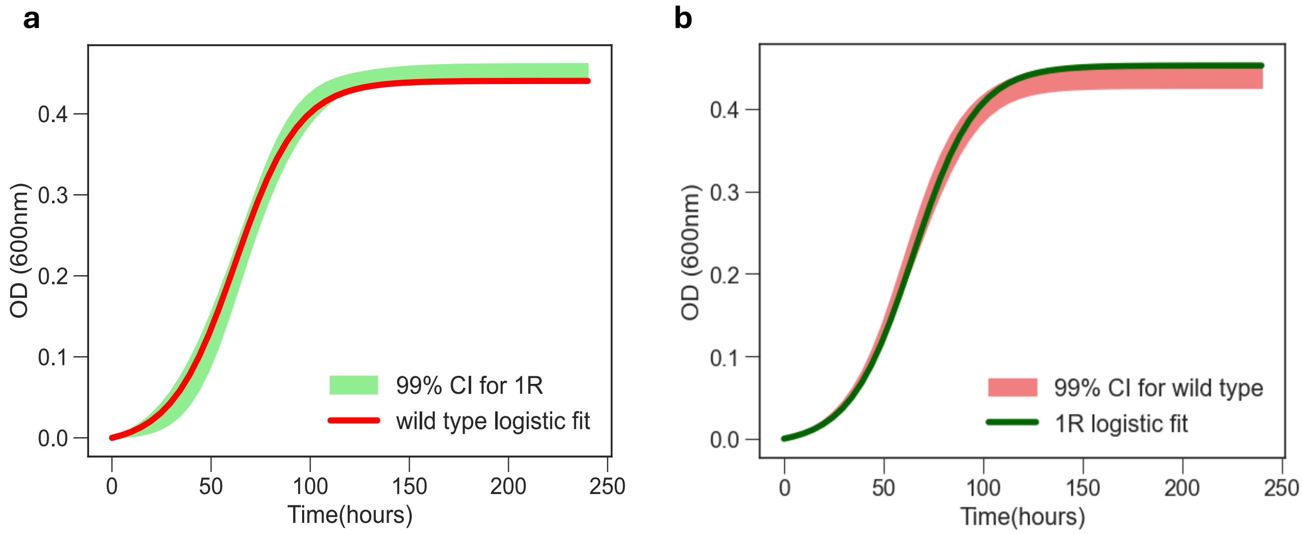


Figure S13: Comparing the growth curves of the wild type *Sal ruber M1* and pseudolysogen 1R from Figure 3a. **a**, Growth curve of *Sal ruber M1* within the 99 % confidence interval of the growth curve of pseudolysogen 1R. **b**, Growth curve of pseudolysogen 1R within the 99 % confidence interval of the growth curve of *Sal ruber M1*.

#### Alternative model: Viral Decision

The main 'split inheritance' model used in this study suggests that the parent pseudolysogen cell has $k+1$ splits to distribute $k$ viral genomes into the daughter cells. This process produces a daughter cell with $i$ viral genomes with a probability of $2/(k+1)$. However it could be argued that instead of each split being completely random and independent of the viral genomes, perhaps each viral genome makes a 50/50 decision to go to one or the other daughter cell. This new process would produce a daughter cell with $i$ viral genomes with a probability of $2\binom{k}{i}(\frac{1}{2})^k$. The model is described by the following system:

$$\dot{R} = -\epsilon\psi(R)(S + \frac{1}{1-\alpha}(\sum_{i=1}^{k} P_e^{[i]} + \sum_{i=1}^{k} P_f^{[i]}))$$

$$\dot{S} = \overbrace{\psi(R)S}^{\text{growth/consumption}} - \overbrace{\phi SV}^{\text{infection}} + \overbrace{2\frac{\psi(R)}{1-\alpha}(\binom{1}{0}(\frac{1}{2})^1 P_e^{[1]} + ...\binom{k}{0}(\frac{1}{2})^k P_e^{[k]} + \binom{1}{0}(\frac{1}{2})^1 P_f^{[1]} + ...\binom{k}{0}(\frac{1}{2})^k P_f^{[k]})}^{\text{reproductive gain}}$$

$$\dot{P_e^{[1]}} = \overbrace{\phi SV}^{\text{infection}} - \overbrace{\phi P_e^{[1]} V}^{\text{superinfection}} - \overbrace{\lambda_1 P_e^{[1]}}^{\text{transition to full formation}} - \overbrace{\frac{\psi(R)}{1-\alpha} P_e^{[1]}}^{\text{reproductive loss}} + \overbrace{2\frac{\psi(R)}{1-\alpha}(\binom{1}{1}(\frac{1}{2})^1 P_e^{[1]} + ...\binom{k}{1}(\frac{1}{2})^k P_e^{[k]})}^{\text{reproductive gain}}$$

$$\dot{P_e^{[2]}} = \overbrace{\phi P_e^{[1]} V}^{\text{superinfection in}} - \overbrace{\phi P_e^{[2]} V}^{\text{superinfection out}} - \lambda_2 P_e^{[2]} - \frac{\psi(R)}{1-\alpha} P_e^{[2]} + 2\frac{\psi(R)}{1-\alpha}(\binom{2}{2}(\frac{1}{2})^1 P_e^{[1]} + ...\binom{k}{2}(\frac{1}{2})^k P_e^{[k]})$$

$$\vdots$$

$$\dot{P_e^{[k']}} = \phi P_e^{[k'-1]} V - \phi P_e^{[k']} V - \lambda'_k P_e^{[k']} - \frac{\psi(R)}{1-\alpha} P_e^{[k']} + 2\frac{\psi(R)}{1-\alpha}(\sum_{i=k'}^{k} \binom{k}{k'}(\frac{1}{2})^k P_e^{[k']})$$

$$\vdots$$

$$\dot{P_e^{[k]}} = \phi P_e^{[k-1]} V - \lambda_k P_e^{[k]} - \frac{\psi(R)}{1-\alpha} P_e^{[k]} + 2\frac{\psi(R)}{1-\alpha}(\frac{1}{2})^k P_e^{[k]}$$

$$\dot{P_f^{[1]}} = \overbrace{\lambda_1 P_e^{[1]}}^{\text{transition to full formation}} - \overbrace{\frac{\psi(R)}{1-\alpha} P_f^{[1]}}^{\text{reproductive loss}} + \overbrace{2\frac{\psi(R)}{1-\alpha}(\binom{1}{1}(\frac{1}{2})^1 P_f^{[1]} + ...\binom{k}{1}(\frac{1}{2})^k P_f^{[k]})}^{\text{reproductive gain}} - \overbrace{\eta\gamma P_f^{[1]}}^{\text{lysis}}$$

$$\dot{P_f^{[2]}} = \lambda_2 P_e^{[2]} - \frac{\psi(R)}{1-\alpha} P_f^{[2]} + 2\frac{\psi(R)}{1-\alpha}(\binom{2}{2}(\frac{1}{2})^1 P_f^{[1]} + ...\binom{k}{2}(\frac{1}{2})^k P_f^{[k]}) - \eta\gamma P_f^{[2]}$$

$$\vdots$$

$$\dot{P_f^{[k']}} = \lambda_2 P_e^{[k']} - \frac{\psi(R)}{1-\alpha} P_f^{[k']} + 2\frac{\psi(R)}{1-\alpha}(\sum_{i=k'}^{k} \binom{k}{k'}(\frac{1}{2})^k P_f^{[k']}) - \eta\gamma P_f^{[k']}$$

$$\dot{P_f^{[k]}} = \lambda_k P_e^{[k]} - \frac{\psi(R)}{1-\alpha} P_f^{[k]} + 2\frac{\psi(R)}{1-\alpha}(\frac{1}{2})^k P_f^{[k]} - \eta\gamma P_f^{[k]}$$

$$\dot{V} = \overbrace{\beta\gamma\eta \sum_{i=1}^{k} P_f^{[i]}}^{\text{burst}} - \overbrace{\phi(S + \sum_{i=1}^{k} P_e^{[i]})V}^{\text{total infection}}.$$

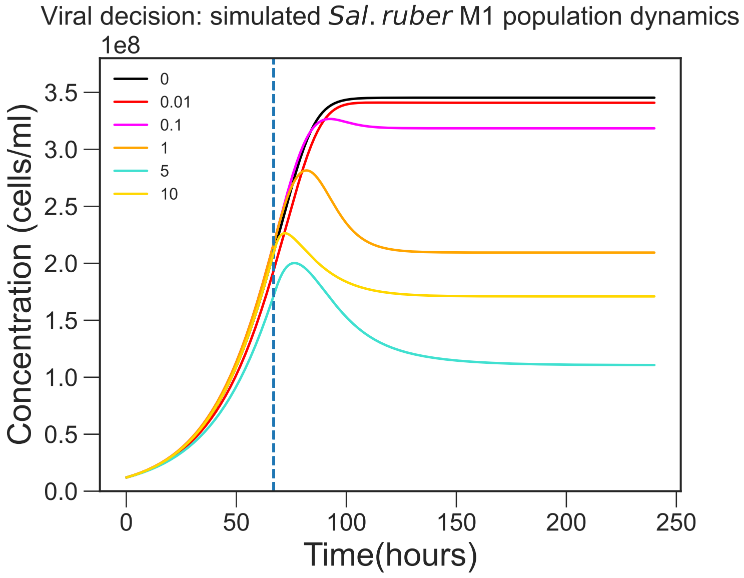


Figure S14: Population dynamics of M1 host given the exact same parameters and conditions as in Figure 5a. M1 host population recovers to a higher population when infected at MOI 10 compared to MOI 5, highlighting the same non-monotonic relationship between viral levels and population-level lysis as seen with split inheritance. However, the MOI 10 crashes quicker upon infection than observed in Figure 5a, highlighting a lack of the persistent transient growth that is a hallmark feature of the dynamics observed experimentally in Figure 3c.

### Episomal Viral Replication

While viral replication in lysogeny generally requires integration of the viral genome into the host chromosome, some viruses such as phage Ff and Pf1, have the ability to replicate episomally without any integration (24) akin to multicopy plasmids (43). As we cannot say with certainty that the EM1 viral genome does not replicate inside the M1 host cell, it may very well be possible that similar episomal replication is occurring within our system. To account for this, we introduce a model that allows for viral replication independent of host cell division as follows:

$$\dot{R} = -\epsilon\psi(R)\Big(S + \frac{1}{1-\alpha}\big(\sum_{i=1}^{k} P_e^{[i]} + \sum_{i=1}^{k} P_f^{[i]}\big)\Big)$$

$$\dot{S} = \overbrace{\psi(R)S}^{\text{growth/consumption}} - \overbrace{\phi SV}^{\text{infection}} + \overbrace{2\frac{\psi(R)}{1-\alpha} z\Big(\big(\frac{1}{1+1}P_e^{[1]} + \frac{1}{2+1}P_e^{[2]}.....\frac{1}{k+1}P_e^{[k]}\big) + \big(\frac{1}{1+1}P_f^{[1]} + \frac{1}{2+1}P_f^{[2]}.....\frac{1}{k+1}P_f^{[k]}\big)\Big)}^{\text{reproductive gain from split inheritance}}$$

$$\dot{P_e^{[1]}} = \overbrace{\phi SV}^{\text{infection}} - \overbrace{\phi P_e^{[1]}V}^{\text{superinfection}} - \overbrace{\lambda_1 P_e^{[1]}}^{\text{transition to full formation}} - \overbrace{\frac{\psi(R)}{1-\alpha}P_e^{[1]}}^{\text{reproductive loss}} + 2\frac{\psi(R)}{1-\alpha}\Big(z\sum_{i=1}^{k}\frac{1}{i+1}P_e^{[i]}\Big) + \overbrace{(1-z)P_e^{[1]}}^{\text{episomal replication}}\Big)$$

$$\dot{P_e^{[k']}} = \phi P_e^{[k'-1]}V - \phi P_e^{[k']}V - \lambda'_k P_e^{[k']} - \frac{\psi(R)}{1-\alpha}P_e^{[k']} + 2\frac{\psi(R)}{1-\alpha}\Big(z\sum_{i=k'}^{k}\frac{1}{i+1}P_e^{[i]} + \overbrace{(1-z)P_e^{[k']}}^{\text{episomal replication}}\Big)$$

$$\vdots$$

$$\dot{P_e^{[k]}} = \phi P_e^{[k-1]}V - \lambda_k P_e^{[k]} - \frac{\psi(R)}{1-\alpha}P_e^{[k]} + 2\frac{\psi(R)}{1-\alpha}\Big(z\frac{1}{k+1}P_e^{[k]} + \overbrace{(1-z)P_e^{[k]}}^{\text{episomal replication}}\Big)$$

$$\dot{P_f^{[1]}} = \overbrace{\lambda_1 P_e^{[1]}}^{\text{transition to full formation}} - \overbrace{\frac{\psi(R)}{1-\alpha}P_f^{[1]}}^{\text{reproductive loss}} + 2\frac{\psi(R)}{1-\alpha}\Big(z\sum_{i=1}^{k}\frac{1}{i+1}P_f^{[i]} + \overbrace{(1-z)P_f^{[1]}}^{\text{episomal replication}}\Big) - \overbrace{\eta\gamma P_f^{[1]}}^{\text{lysis}}$$

$$\vdots$$

$$\dot{P_f^{[k]}} = \lambda_k P_e^{[k]} - \frac{\psi(R)}{1-\alpha}P_f^{[k]} + 2\frac{\psi(R)}{1-\alpha}\Big(z\frac{1}{k+1}P_f^{[k]} + \overbrace{(1-z)P_f^{[k]}}^{\text{episomal replication}}\Big) - \eta\gamma P_f^{[k]}$$

$$\dot{V} = \overbrace{\beta\gamma\eta\sum_{i=1}^{k} P_f^{[i]}}^{\text{burst}} - \overbrace{\phi(S + \sum_{i=1}^{k-1} P_e^{[i]})V}^{\text{total infection}}$$

Where $z$ represents the probability that upon host cell division, the virus does **not** replicate. Therefore the viral genome replicates every $\frac{1}{1-z}$ host doubling generations (if $z = 1$, viral genome never replicates akin to main model). Figure S15 shows how the results change from no viral genome replication (equivalent to Figure 5a in Main Text) to perfectly synchronous replication ($z = 0$) on a continuum. The results shows that the more frequent virus replication is, the less of a lysis induced dip is observed at MOI 5 (and to a lesser degree at MOI 10 as well). The dip and recovery dynamic is a hallmark feature of the MOI experiment (Fig 5a), therefore highly synchronous viral replication that omits this feature as seen in Figure S15d (every 1.25 host generations) and S15e (every host generation) is unlikely. However, the dip and recovery dynamic, as well as the rest of the key qualitative features of the experimental results, are maintained when the model allows for less frequent viral replication as seen in Figure S15b (every 5 generations) and S15c (every 2 generations). This highlights that a degree of extra chromosomal viral replication might be present in the M1-EM1 system while retaining the key non monotonic features of the experiment.

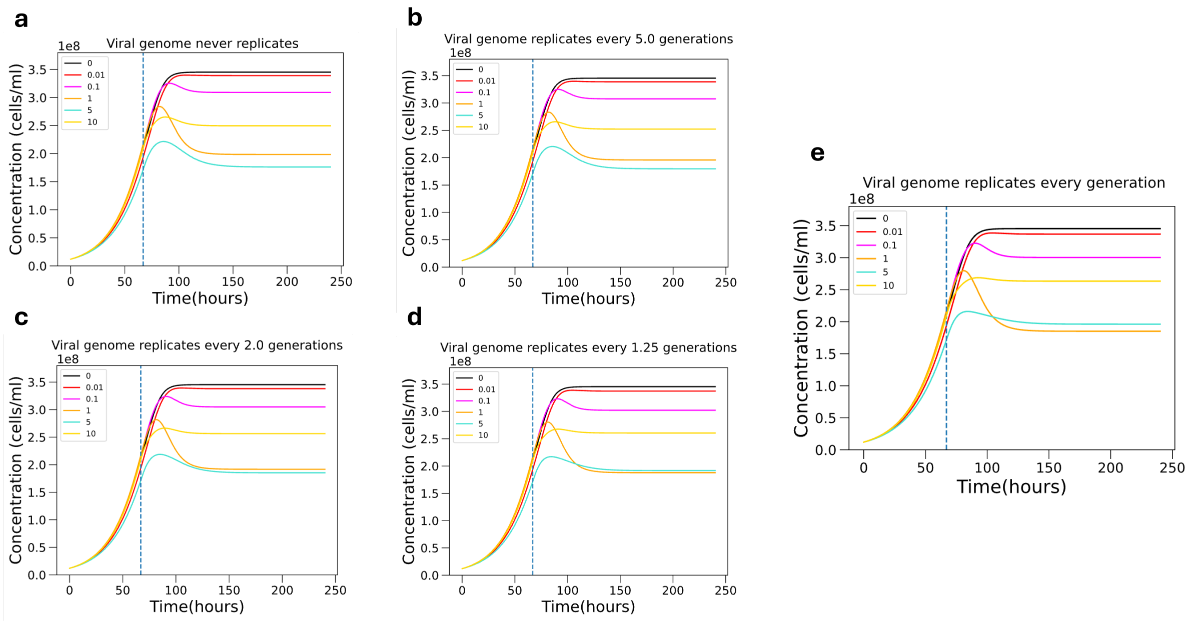


Figure S15: Population dynamics of Sal Ruber M1 upon infection by EM1 in the presence of potential viral replication independent of the host where **a,** $z = 1$, **b,** $z = 4/5$, **c,** $z = 1/2$, **d,** $z = 1/5$, **e,** $z = 0$. Generations are defined in terms of doubling times of the M1 population. Note that Figure S15a is the equivalent of the main model graph seen in Figure 5a in the main text.
